# Red states modulate energy localization between the antenna and trap in PSI

**DOI:** 10.64898/2026.10.01.756128

**Authors:** Marvin Asido, Halima Khatun, Anton Khmelnitskiy, Mike Reppert, Gabriela S Schlau-Cohen, Yuval Mazor

## Abstract

The core antenna of the photosystem I (PSI) complex of oxygenic photosynthesis operates at near perfect quantum efficiency, making it one of the most efficient energy conversation devices in nature. The presence of low energy or red chlorophylls (chls) is nearly universal in this core antenna, and these red chls are known to extend the light harvesting capacity and control the rate of energy migration in PSI. Surprisingly, the significance of the structural position of the red chls in PSI remains unknown. In this study, we introduced mutations that generate specific red sites in the PSI core antenna, consistent with those observed in natural species. Using CryoEM, we characterized the structural changes at these red sites. Using transient absorption spectroscopy, we measured the effects of red chlorophyll at specific sites on energy transfer and trapping in PSI to identify their functional significance. While one site funneled energy to the reaction center, the other slowed equilibration in the antenna. These opposing effects indicate different functions are associated with the different red chlorophyll location, reconciling previous seemingly incompatible suggestions as to their functional role.

## Introduction

Light harvesting in oxygenic photosynthesis relies on two distinct photosystems. Photosynthetic organisms harness photon energy to oxidize water (photosystem II, PSII) and produce NADPH and ATP for carbon fixation (photosystem I, PSI) (Blankenship R, 2021). The core structures of both photosystems are highly conserved among cyanobacteria, which appeared approximately 3 billion years ago, and all photosynthesizing eukaryotes, which diverged from cyanobacteria approximately 1.5 billion years ago, showing that photosystem evolution is highly constrained (Ben-Shem et al., 2003; Jordan et al., 2001). Both photosystems use a dense array of chlorophyll A (chl) molecules as a core antenna, and as part of their internal electron transport chains. Photosystem I is the more extreme example of this, with around 90 chl molecules in its core antennae amounting to a concentration of ∼0.5 M (Croce & Van Amerongen, 2013). The chl molecules transport and convert photoenergy with near perfect quantum efficiency, making it the most efficient photochemical devices in nature. This high efficiency also minimizes side reactions which contributes to the high resilience of PSI, with regards to both high temperatures and minimizing the formation of damaging oxygen species. Each of the 90 chl that make up the core antenna in PSI has a transition energy uniquely tuned by its environment, known as a site energy. In addition, close contacts with neighboring chl result in excitonic coupling between the chls that further modify the observed energies. The relationship between the energetic landscape of these core antenna chls and other properties of PSI is not clear.

One surprising property of many PSI complexes is the presence of a subset of low energy or red chl in their core antenna. These are defined as chl with excitation energy lower than that of P700, the stable electron donor in PSI. The existence of red chls implies that certain steps in the excitation energy transfer processes in PSI occur energetically uphill so must be thermally activated (Giera et al., 2018; Hayes et al., 2000; Hsin et al., 2004; Shubin et al., 1992; Zazubovich et al., 2002). As a result, the red chls are the source of relatively slow transfer steps that limit the overall rate of energy transport through the PSI core, ranging from around 20 ps in PSI systems with low red chl content, to as high as 50 ps in systems with high red chl content, such as Spirulina (Gobets, Van Stokkum, et al., 2001). Despite the clear effects of the red chls, the actual function is not as obvious. Extending the absorption of PSI into the red region of the spectrum is one benefit (H.-W. Trissl, 1993; Rivadossi et al., 1999). However other functions such as photoprotection have been suggested in the past (Schlodder et al., 2011). In cyanobacteria, estimations on the number of red chl based on various spectral measurements differ between species (Karapetyan et al., 2014), however whether these differences represent enhancement of conserved low energy sites or from random changes across the PSI antenna is still undermined. At lower temperatures, in most species the emission spectrum from PSI is dominated by red chls, i.e., they are the terminal emitters of the core antennae.

In contrast to the large body of spectroscopical observations that have accumulated on red chls in PSI (Croce et al., 1998; Giera et al., 2018; Gobets et al., 1994, 2003; Gobets, Van Stokkum, et al., 2001; Hayes et al., 2000; Hsin et al., 2004; Shubin et al., 1991), structural identification of the chls responsible for red sites is lacking. First, although there are quite a few structures of PSI, determining site energies is challenging. Second, optical data which can be used to phenomenologically determine site energies, does not carry sufficient information to identify a unique set of values. Third, the relatively small number of red chl (<10) compared to the overall amount of chl within the complex (> 90) complicates a clear temporal and spectral separation of the respective contributions in a (time-resolved) spectroscopic experiment. The inability to connect spectroscopic observables to their spatial origin has been a major barrier to identifying the function of the red chl.

In this study, we introduced mutations that generate specific red sites in the PSI core antenna, consistent with those observed in natural species. Using CryoEM, we characterized the structural changes at these red sites. Using transient absorption spectroscopy, we measured the effects of red chl at specific sites on energy transfer and trapping in PSI.

## Experimental/Methods

### Growth condition

Synechocystis wt and mutant strains were grown on BG11 plates in a 30 °C room under continuous white light (∼40 μmol⋅m^−2^⋅s^−1^ photons) for two weeks. When the growth of cyanobacteria was sufficiently dense, they were removed from the plate to a sterile flask which contained 50 ml liquid BG11 media supplemented with 5mM glucose, 6 μg/ml Ferric ammonium citrate and 10 μg/ml kanamycin (for mutant strains) at the same light and temperature condition. After two weeks the cultures were transferred to 500 ml Liquid BG11 media. After two weeks the cultures were transferred to 10 L glass bottles for another two weeks of growth.

### Preparation of thylakoid membrane

Cells were spun down using centrifugation (LYNX 6000, Thermo Scientific) in a F9-6×1000 LEX rotor in for 5 minutes at 9000 rpm. Then they were washed with ∼250 ml STN1 buffer (30mM Tricine-NaOH pH 8, 15 mM NaCl and 0.4 M Sucrose) and spun down again in F20-12×50 LEX rotor for 10 minutes at 12,000 rpm. The cells were then resuspended with ∼50 ml STN1 solution and were broken using a cell disrupter (constant systems) at 30,000 psi for 3 cycles. The lysate was spun down in F20-12×50 LEX rotor for 5 minutes at 12,000 rpm and the supernatant was collected. Membranes were pelleted using ultracentrifugation (Ti70 rotor) at 45,000 rpm for 2 hours at 4 °C. Pellets were collected and resuspend in STN2 (30mM Tricine-NaOH pH 8, 150 mM NaCl and 0.4 M sucrose) solution. Following 15 minutes incubation on ice the membranes were sedimented again using an ultracentrifuge (Ti70 rotor) at 45,000 rpm for 2 hours at 4 °C. The membrane pellet was collected and resuspend in ∼15 ml STN1 solution. Concentration of chl was then measured according to Arnon (Arnon, 1949). The resuspended pellet was then stored in −80 °C.

### Isolation of the PSI complex from cyanobacteria

Thylakoid membranes from wt and mutants were solubilized with n-dodecyl-beta-maltoside (DDM, Glycon). DDM was added to achieve a ratio of 15:1 DDM to chl mass. The suspension was mixed manually by the inversion of the tube a few times and incubation for 30 minutes at 4 °C. To remove insoluble materials, thylakoid membranes were then spun down on ultracentrifuge (Ti70 rotor) at 45,000 rpm for 30 minutes at 4 °C. The supernatant was loaded into the diethylaminoethyl cellulose (DEAE) gravity flow column (Toyopearl C650) followed by the addition of a linear gradient of NaCl (15-250 mM NaCl) in 30 mM Tricine-NaOH at pH 8.0 and, 0.1% DDM to elute the complexes. The dark green fractions that correspond to PSI were then collected followed by the addition of PEG3350 (Hampton research) to a final concentration of 8%. The PSI trimer was then precipitated using F20-12×50 LEX rotor for 5 min at 5000 rpm. The green precipitation was resuspended in 30 mM Tricine-NaOH pH 8, 0.05% DDM, and 75 mM NaCl and was loaded onto 10%-30% sucrose gradients in a SW40 rotor and centrifuged at 36,000 rpm for 12-16 hours. The green band was then collected from the sucrose gradient.

### Sample preparation for single-particle cryo-EM analysis

The second green band from the sucrose gradient was collected, the NaCl concentration was brought to 250 mM and PEG3350 was added to 8% to precipitate the complex. Following precipitation (5 min, 14,000 rpm) the complex was resuspended in 30 mM Tricine pH8, 75 mM NaCl, 0.002% DM and a 3-µl drop of the PSI complex (concentration 3 - 5 mg/ml chl) was applied on to a holey carbon grid (C-flat 1.2/1.3 Cu 300-mesh grids, Protochips) after soaking the grid in buffer. The sample was vitrified by flash-plunging the grid into liquid ethane using an automated plunge freezer, a Vitrobot Mark IV (ThermoFisher/FEI) with a blotting time of 3-6 s and blotting force of −1. The grids were stored in liquid nitrogen before data acquisition.

### Ultrafast transient absorption spectroscopy

Femtosecond transient absorption spectra were collected using a commercial femtosecond transient absorption spectrometer (Helios Fire, Ultrafast System). The 800 nm fundamental pulses were generated from a Ti: Sapphire laser system (Astrella, Coherent Inc., <100 fs, 7 mJ/pulse, 1 kHz repetition rate) and were split into two beams. One beam was used to generate a white-light supercontinuum (WL) generated by focusing the beam on a 2 mm sapphire plate inside Helios Fire as the probe. The second beam was used to generate 425 nm pump pulses via an optical parametric amplifier (OPerA Solo, Coherent Inc.). The pump power was then reduced to excite no more than 1% of PSI complexes using a neutral density filter. The pump pulses were depolarized before focusing on the sample. PSI samples reduced with sodium ascorbate (40 mM) and phenazine methosulfate (PMS, 40 μM) were loaded in a home-built spinning wheel cuvette with a final OD_680_ of ∼0.6 in an optical path length of 1.2 mm. All measurements were performed at room temperature.

### Lifetime Distribution Analysis

The time-resolved spectroscopic data was analyzed using OPTIMUS (Slavov et al., 2015), an analysis software available free-of-charge at www.optimusfit.org. The experimental data was objected to the (quasi) model-free lifetime distribution analysis (LDA) (Croce et al., 2001; Slavov et al., 2015), using Tikhonov regularization with 100 regularization factors and 200 evenly distributed lifetime components. The best fit is obtained by using the L-curve criterion, which represents the minimum smoothing factor vs residual norm. As a result of the LDA, the lifetime distribution of the individual state transitions was obtained, which is graphically represented in the corresponding lifetime distribution map (LDM). Together with the 2D representation of the transient data, the LDMs reveal complex kinetic behavior (e.g. multiphasic decays) and spectral shifts and allow a good first approximation of the expected lifetime components. The sign of the lifetime distribution amplitude can correspond to a decay or a rise of a species. A positive amplitude in the lifetime distribution corresponds to a decaying species (transient signal is positive) or a population of a new species (transient signal is negative). A negative amplitude in the lifetime distribution corresponds to a decaying species (transient signal is negative) or a population of a new species (transient signal is positive). Note that when using specific lifetimes in the text, we refer to the center of each distribution.

### Global target analysis

A (bio)physical model was obtained by global target analysis (GTA), which connects species associated difference spectra (SADS) with each kinetic component of the model(Beechem et al., 1985; Müller et al., 2003; Roelofs et al., 1992; van Stokkum et al., 2004). The ‘quality’ of the SADS is judged based on knowledge of the spectral features of the expected intermediate species and their temporal sequence. In the case of PSI, five species were considered which are: excited antenna (core antenna), excited red chls (red*), excited special pair (P700*) and two charge separated states (P700^+^/A_0_ and P700^+^/A_1_)(Müller et al., 2003). To build the model, we used kinetic information obtained by the lifetime ranges in the LDMs, the analysis of the ΔΔ datasets and the time-resolved fluorescence datasets. The lifetimes were used as starting parameters for the τ_1_, τ_2_, τ_4_ and τ_5_ components in the GTA and were kept soft-fixed within a margin of 5-10%. The τ_3_ component, which falls within the most complex temporal range, was kept free. Out of all the kinetic connectivities tested within a 5-compartment model, only a 5-compartment model with equilibria between S1/S2 and S2/S3 leads to reasonable SADS and lifetimes within the margin of the set parameters for all samples.

### Biased hexagonal antenna arrays

Excitation migration, simplified to a single hope migration between nodes, was simulated on a hexagonal array containing 91 nodes. Rates were used to define a central node representing the RC, antenna nodes and an exponential term was used to introduce biased rate to one antenna node representing the low energy site (see **Figure S19**). A 1000 runs initiated from a random node were used to evaluate trapping time for each energy well location. Simulation were performed using an inhouse python script available from the authors upon request.

### CryoEM data collection

Micrographs were acquired using a Krios I (Thermo Fisher Scientific) operated at 300 kV and a magnification of 165000x. Images were recorded on a K3 (Gatan Inc.) equipped with a SelectrisX energy filter with a slit width of 20 eV. The targeted defocus range was varied from −0.8 to −2.5 µm using the EPU software (Thermo Fisher). Data processing was carried out using CryoSPARC 4.7m (Punjani et al., 2017). Full description of the data processing pipeline and descriptive statistics are available in Figures S2-5 and table S1.

### Model building and refinement

The initial PSI model was taken from the RedA PSI structure (PDB: 6UZV) (Toporik et al., 2020). The model was docked into the map using UCSF ChimeraX(Pettersen et al., 2021). The RedB mutation was introduced using Coot (Emsley et al., 2010) which was also used to manually adjust the model into the map. The final model was refined against the Cryo-EM density map using phenix.real_space_refine (Liebschner et al., 2019). Final model statistics are shown in Table S1. UCSF ChimeraX was used to generate all structural images (Meng et al., 2023).

### Superposing different PSI models and measurement

Superposing and (Round Mean Sqaure Distance) RMSD calculation were done with ChimeraX. Maps were superposed manually first and then using the fit map tool in chimerax until the best match was obtained. All maps were superposed on the RedAB map. Models were first superposed on the RedAB structure using the matchmaker utility in chimerax using chain B (PsaB) for all the models. After this alignment, a map (at a resolution of 3Å) was calculated from the PsaA and PsaB subunits of RedAB and the models were fitted to this map using the fitmap command in Chimerax. This procedure was chosen to consider the ligands in the alignment process itself.

## Results

### Structural analysis of the RedA site

Structural knowledge on the configuration of red sites is limited, particularly in cyanobacteria. To identify the key characteristics, we determined the structure of the RedAB PSI trimer from *Synechocystis* PCC 6803 (6803). The WT PSI trimer from 6803 (wt 6803) contains a small number of red sites. In the RedAB PSI trimer from 6803, two structural changes were introduced in the PsaB PSI subunit, both naturally occurring in the *Thermosynechococcus vestitus* (TV) PSI complexes. These changes were previously shown to introduce red states into the 6803 PSI (Khmelnitskiy et al., 2020; Toporik et al., 2020).

We used two models fitted into cryoEM maps to describe the structure of the RedAB PSI **(figure s2** & **s3** and **table s1)**. In one, employing C3 symmetry (via C3 expansion of the data, followed by focused refinement, **figure s4**) resulted in an overall resolution of 1.76 Å and represents the average PSI monomer. In the other, reconstruction of the PSI trimer map without symmetry consideration resulted in three distinct monomer orientations with a resolution of 1.9 Å (**figure s5)**. Both the RedA and RedB sites were compared to wt 6803 structure, determined from a crystal structure and refined as three independent monomers as well (Malavath et al., 2018). Other cyanobacterial structures with and without mutations at the RedA and RedB sites were used as additional comparison points (Gisriel et al., 2024; Jordan et al., 2001; Li et al., 2022; Toporik et al., 2020).

**Figure 1.A** shows the location of the RedA and RedB mutations in PSI. RedA consists of a loop insertion into PsaB, which results in the binding of chl B33 (absent from wt 6803), while RedB is a histidine to glutamine change in the amino acid coordinating chl B7 (Khmelnitskiy et al., 2020; Toporik et al., 2020). The RedA site is thought to consist of chlB33 along with its neighboring chl, B31 and B32. As expected, the overall configuration of the RedA chl trimer is highly similar across the RedA, RedAB, and wild type TV PSI, as illustrated in **figure 1.B**.

**Figure 1:**
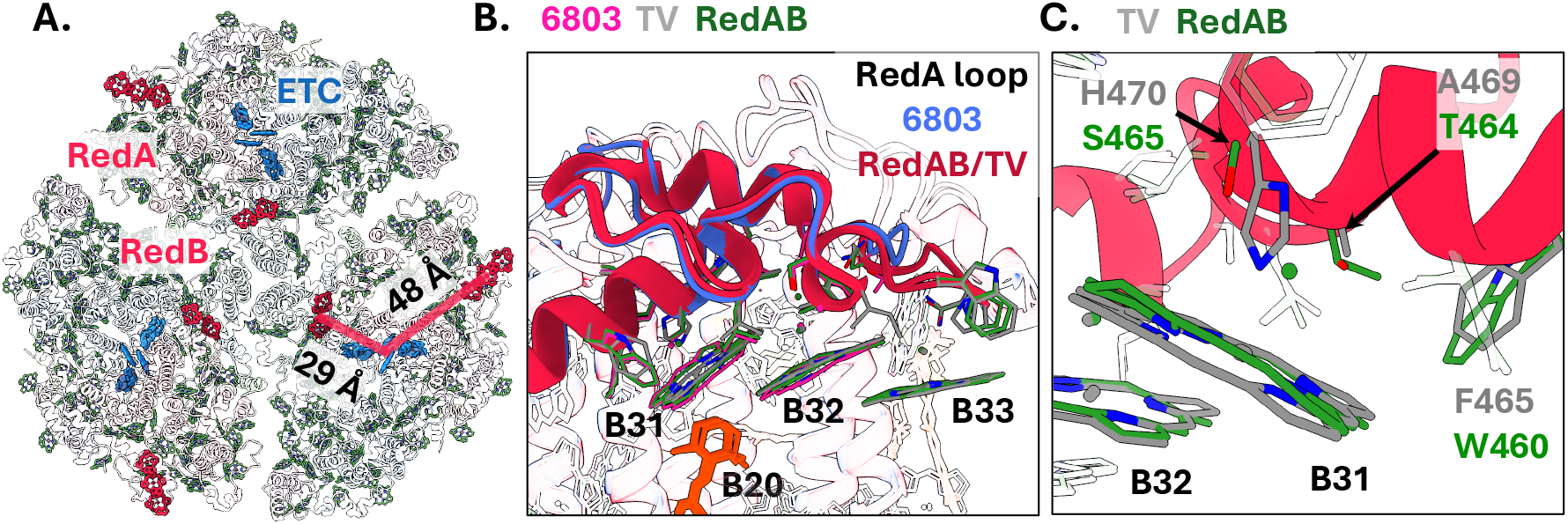
Structure of the RedA site. **A**. Overall view of the RedAB PSI trimer with the two mutation sites indicated in red and the chls of the electron transport chain in blue (the P700 pair is seen as a side view of both chls). The distances between each site and P700 are indicated. **B**. The RedA environment as it appears in two independent structures, TV (PDB 1jb0) and the current RedAB (PDB 9oy9), compared to the non-red containing RedAB background, wild type 6803 (PDB 5oy0). The PsaB loop is show in blue for 6803 and red for the RedAB/TV. Specific amino acids or chls are color coded according to the key at the top of the image. **C**. Amino acid changes separating 6803 RedAB and TV. All amino acids within 5 Ang of the ring atoms of B31-B32-B33 are shown. Common amino acids between RedAB and TV are transparent and the three amino acid changes, all near B31, are shown colored, in green (RedAB) or grey (TV).

The chl site energy can be sensitive to small structural changes (Liguori et al., 2020; Rankelytė et al., 2024; Reiter et al., 2023). However, identifying these differences is challenging because comparisons of the RedA site across species and structures are complicated by the fact that each structure was determined at a different resolution and refined using (potentially) different parameters (Lin et al., 2024). For chl B31, some differences between TV and RedAB exist in the local chl protein pocket, explaining the different conformations observed between RedAB and TV **(figure 1.C**). For chls B32 and B33, small differences are observed between the different structures (**figure 1.B**). Due to the location of the RedA site at the periphery of PSI, where local map resolution decreases, we conclude that for chls B33 and B32 our RedA site structure represents the same conformation as in the TV structure. Chl B32 was identified recently as having one of the lowest site energies in PSI, and this suggests that even in wt 6803 this site constitutes a red chl, with excitonic coupling to B33 probably responsible for the enhanced red nature of this PSI feature in RedAB and TV (Reiter et al., 2023). Calculated electronic coupling values between the RedA chl’s are quite variable (**table s2**) as calculated using the transition electrostatic potential method (Madjet et al., 2006) implemented in the PigmentHunter application (Ahad et al., 2024). This variability reflects the limited resolution of these chls even in the current, relatively high-resolution map.

### Structural analysis of the RedB site

In contrast to the RedA site, the RedB site is located close to the center of the PSI trimer, where the local resolution of the map is high, making small conformational differences significant. The RedB site consist of the chl dimer B7 and A32 (Jordan et al., 2001; Khmelnitskiy et al., 2020). Examining the chl B7 conformation across several structures revealed an unexpected level of structural heterogeneity, with different structures from wt 6803 showing different configurations (**figure 2.A**). To confirm that this heterogeneity does not reflect differences in the modeling approach used to determine the different structures, the maps and models from the different PSI structures were superposed on an entire PSI monomer and are shown around chl B7 in **figures s6-10**. This comparison confirmed that the conformation of chl B7 is highly similar between the wt 6803 (5oy0) structure and RedAB when maps are compared visually and by RMSD values across the chl ring atoms (**figure s6-10)**.

**Figure 2:**
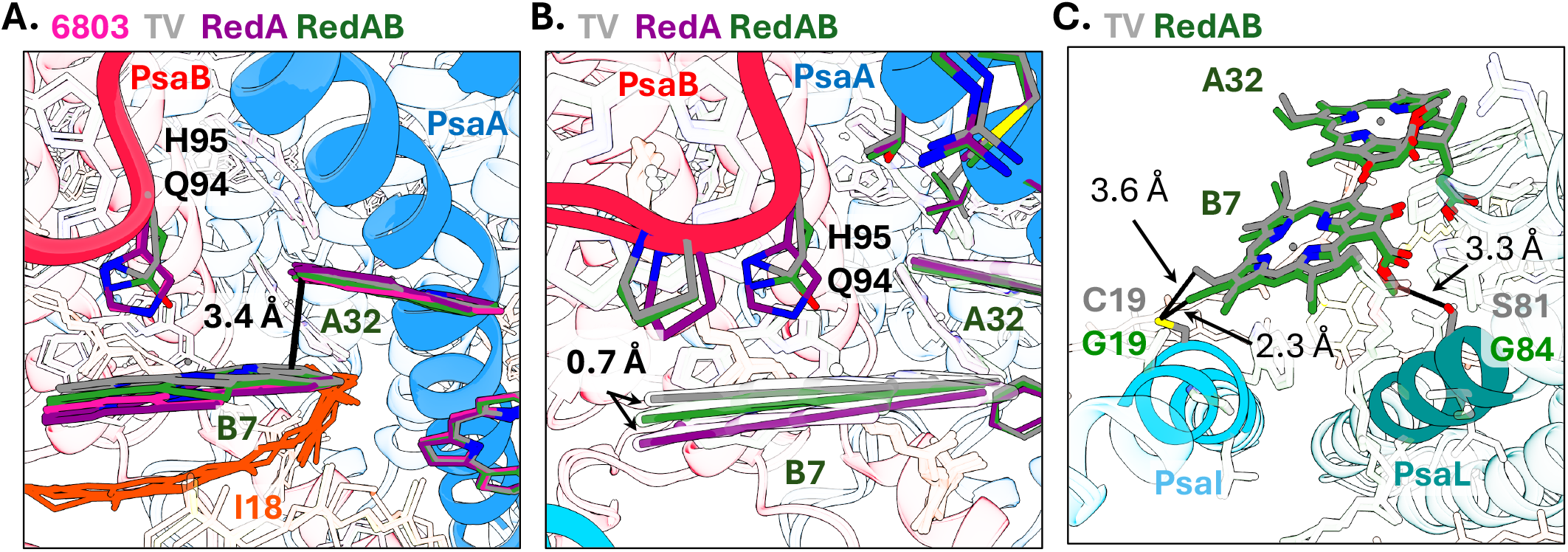
Structure of the RedB site. **A**. The RedB site as it appears in two independent red shifted (glutamine coordinated) structures, TV (PDB 1jb0, 7fix) and RedAB PSI (PDB 9oz3), compared to two, blue shifted (histidine coordinated) structures, wt 6803 (PDB 5oy0) and RedA PSI from 6803 (PDB 6uzv). PsaB (red), PsaA (blue) and PsaI (cyan) are shown in cartoon. The B7 coordinating side chains are shown as sticks, Beta carotenes in orange. Chls and side chains are colored according to key on top of each image, the general similarity between the 6803 wt B7 (in pink) and the RedAB B7 rings (in green) can be seen from this side view. **B**. The Qy dipole of chls B7 and A32 (shown in part) were visualized as lines drawn across the NB and ND chl atoms, the positional changes between RedA (in purple, approximating the wt position), RedAB (green) and TV (grey) are summed by a one sided 0.7 Å movement in the dipole line, with approximately similar distance separating the RedB B7 and the TV B7 positions. **C**. Two remaining differences in the B7 environment, both occupied by glycine residues in 6803 and contain cysteine or serine residues in TV, likely account for the remaining structural differences in the RedB site configuration.

**Figure 2.B** visualizes the dipole orientation from the configuration of chl B7 in the RedB site as compared between wt 6803 and TV, which contains a histidine coordination vs glutamine coordination B7 (blue vs red configuration respectively). The dipole orientation of the chl B7 in RedB is found to be in an intermediate position, between 6803 and TV. We calculated the coupling values between chls B7-A32 and A31 (**table s3**). These values indicate, as previously shown, strong coupling of chls B7 and A32 ranging from 118 to 130 cm^-1^. The strongest coupling values were found for the structures containing the RedB site (RedAB and TV), which are glutamine coordinated. However, the differences in coupling values are small and are comparable, for example, to those observed using different methodologies to compute excited states properties (Madjet et al., 2006).

The remaining differences between the sites are visualized in **figure 2.C**. Two glycine residues near chl B7 in wt 6803 (located on the PsaI and PsaL subunits) are replaced by either a cysteine or a serine residue in TV. These changes can both displace chl B7 due to steric considerations or modify the electrostatic environment to some extent, and both factors could contribute to additional spectroscopic differences between the two species.

To estimate the effect of the histidine to glutamine substitution on chl ring distances at the RedB site, we examined several PSI models with either histidine or glutamine coordinating chl B7. In the group of PSI models with histidine coordination, the average ring separation between B7 and A32 in this group is 3.93±0.12 Å, interestingly, the wt 6803 (5oy0) values fall outside this SD interval despite being included in this group. In the group of PSI models with glutamine coordination, the average ring separation is ∼0.2 Å shorter (3.75±0.04 Å, **figure 3.B**). Analysis of the four known red sites, P700 and three red sites in the eukaryotic LHCI, shows an average ring separation of 3.68±0.10 Å (**figure 3.C**). Our conclusion is that the distances between chl B7 and A32 in RedAB are consistent with a red chl pair. The unusual shorter ring separation of 3.8 A observed exclusively in one wt 6803 structure (5oy0) as compared to other structures of the same species. Given the small mutation induced changes we observed (0.2 – 0.5 Å), this “red” conformation may be attainable due to pressure from crystal contacts or dehydration effects (Eyal et al., 2005; Jacobson et al., 2002).

**Figure 3:**
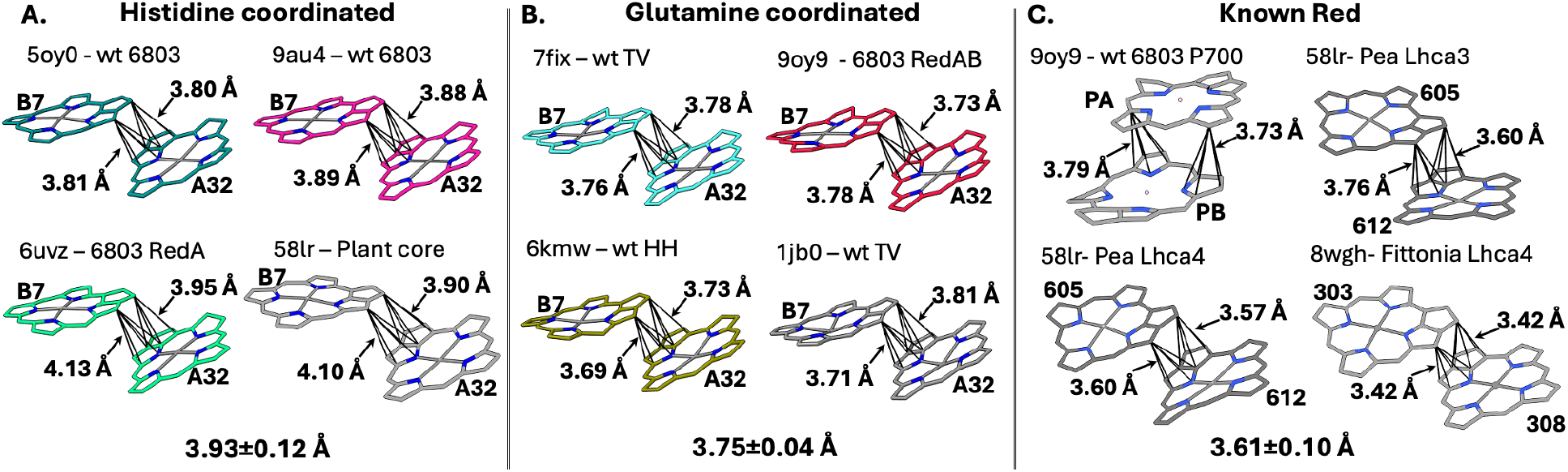
Distances between chl ring systems across different sites. For each chl pair we measured the distances from the closest carbon atom in one ring (out of the five chls rings) to all atoms in the closest ring on the opposite chl, and the average distance is displayed. At the bottom of each section is the average and standard deviation values for each group. **A**. Four structures where chl B7 is coordinated by a histidine sidechain. **B**. Four structures where chl B7 is glutamine coordinated. **C**. Four samples from known red sites. P700 from 6803 and three known red sites from the plant LHCI antenna.

Overall, the histidine to glutamine change in the RedB site results in small structural changes of ∼0.15 Å, which are smaller than those observed in analogous situations. For example, in lhca3 and lhca4 from plants where chl coordination is modified from asparagine (red-shifted) to histidine (blue-shift) there is a structural change of 0.7 Å (Capaldi et al., 2025; Mazor et al., 2015). Collectively, these data suggest that a small ring separation is a rough proxy for a red shift. Surprisingly, our survey of the RedB sites across different PSI structure from 6803 suggest that under certain circumstances a histidine coordinated B7 chl can be in a similar conformation to that observed under glutamine coordination, potentially indicating an ability to adopt a red chl conformation.

### Structural variations across the PSI trimer

The observation that the RedB site displays some heterogeneity across different PSI models prompted us to explore the differences across the PSI monomers in our C1 refined structural model. We calculated the RMSD between the three PSI monomer pairings. The results show that most of the differences are seen in PsaL subunit which coordinate trimer assembly (**figure s11**). High RMSD values correlated with (and potentially explained by) low map resolution are observed where PSI subunits interact with the surrounding membrane, this is seen close to the PsaK, F and J subunits and in the membrane bordering regions between the three PSI monomers (comparing **figure s12** and **figure s11** with **figure s5**). In contrast, the highest RMSD values are observed at the C terminus of the PsaL subunit, which was determined at a high local resolution (∼2 Å), supporting the notion that it represents a different distinct conformation (**figure s11.B and C**). To extend these observations to pigments, we determined the average RMSD value across all chl (measured over the chl ring atoms) and Carotenes. The results are shown in **figure s12** for the entire PSI trimer and follow the general trends observed for the protein subunits. We plotted the average chl RMSD values Vs either local resolution or the model b-factors in **figure s13**, both plots show a moderate correlation (∼0.4 Pearson’s coefficient), with most chl RMSD values fall within 95% confidence intervals. One clear exception is the PsaL chl L1, for which the RMSD values falls outside the model’s SD interval and represent a probable structural change, albeit with currently unknown functional consequences. Examining the RedB site specifically, **figure s11** shows that chl’s B7 and A32 are both determined at very high local resolution, with chl B7 showing medium RMSD values (0.14 Å), somewhat higher than that measured for Chl A32 (0.09 Å). This observation is consistent with the variability we observed in the different structures. For chl B33, part of the RedA chl trimer (**figure s13**), RMSD values are lower than those of chl L1 and close to those of B7, leaving room to the notion that it might have significant conformational variability. This is supported by its high b-factor and low local resolution (**figure s13**).

### Kinetic analysis

To explore the functional effects of the RedA and RedB sites on light harvesting and primary charge separation, we used spectrally-resolved femtosecond transient absorption (TA) spectroscopy to determine the excited-state dynamics in PSI. The relevant spectral window of the TA data for RedA and RedB is plotted (**figure 4.A**, WT and RedAB in **figure s14.A**). The TA maps contain the characteristic difference absorption signatures: i) a ground state bleach (GSB) at the center wavelength of the core antenna chl, which transitions into the redshifted GSB features of the special pair and the red chl; ii) a chl-associated excited state absorption (ESA) at the blue end of the spectral window; and iii) a broad and long-lived absorption feature at the red end of the spectral window, which is a mixture of chl-associated ESA and absorption of the transiently formed radical pairs.^3,4^

**Figure 4:**
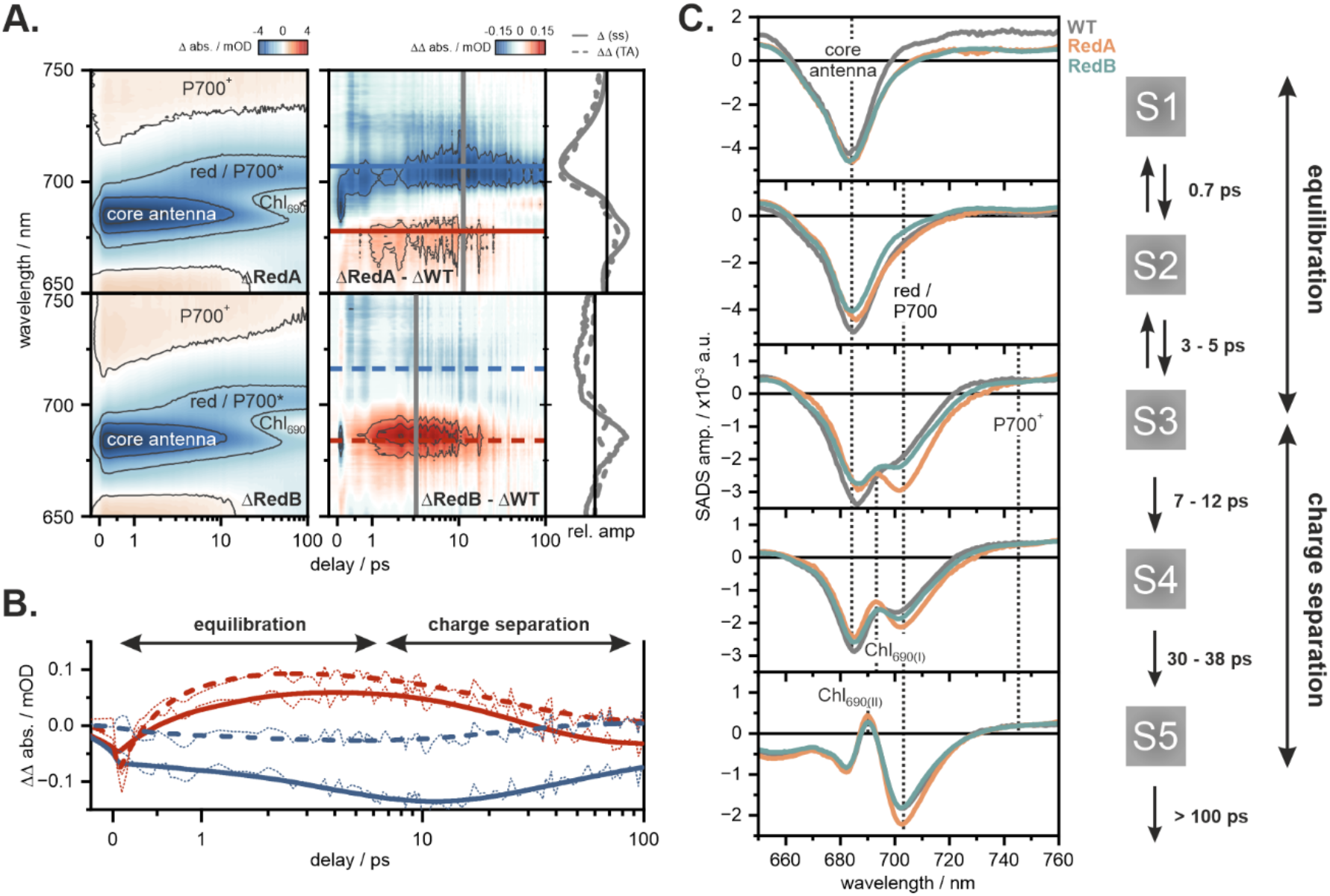
Site-dependent ultrafast dynamics and kinetic model. **A**. Ultrafast transient data of RedA and RedB in the reduced spectral range of 650-750 nm (left) and the corresponding double difference (ΔΔ) spectra after subtraction of the wildtype dataset (right). The spectral cuts (grey, solid line) are overlayed with the inverted and scaled steady state difference (Δ) spectra (grey, dashed lines). **B**. The temporal evolution of the ΔΔ data is extracted by horizontal cuts in the core antenna ΔΔ band (red) and red/P700* ΔΔ band (blue). RedA shows a slow rise of the positive core antenna ΔΔ band and strong accumulation of the negative red/P700* ΔΔ band. In contrast, the core antenna ΔΔ band in RedB rises quickly whilst the red/P700* ΔΔ band is barely accumulated, underlining a significant difference in energy propagation between the core antenna, the respective red sites in RedA and RedB and the PSI ETC. **C**. Compartment model obtained by GTA. The SADS of WT (grey), RedA (orange) and RedB (green) are overlayed. The vertical lines mark spectral features which are characteristic for the individual species. Compartments S1 and S2 represent energy equilibration, S4 and S5 represent charge separation. Compartment S3 contains features of both processes. The RedAB spectra are included in **figure s14.C**.

### Normalization and double differences

We isolated the red site-specific dynamics by subtracting the wildtype data from the data for each mutant (double difference, ΔΔ, **figure 4.A, B** and **figure s14.A**). The ΔΔ spectra reveal a doublet signature with a positive (around ∼680 nm, mostly antenna chl) and negative (around ∼705-715 nm, red chl) contribution, reflecting a reduction in the blue component (∼680 nm) and an increase in the red component (∼705-715 nm) in the mutants relative to the wildtype consistent with the red-shift as observed in the steady state Δ spectra (Abs_RedA,B,AB_ – Abs_WT_, **figure 4.A**, far right, solid gray line; **figure s15** and (Khmelnitskiy et al., 2020; Toporik et al., 2020)). The temporal evolution of the ΔΔ bands was sufficiently described by a set of four lifetimes τ_1_-τ_4_ (**table s5**). Noteworthy are the lifetimes τ_2_ and τ_3_, which in the case of RedB (**figure 4.B, dashed lines**) indicate a fast population and fast depopulation of the red component, whereas in the case of RedA (**figure 4.B, solid lines**) indicate the opposite. In the case of RedAB we observe a fast population and slower depopulation of the red states (**figure s14.A** and **B**). These observations already mean qualitatively that site A and site B have different effects on the kinetics.

### Global Target Analysis and Kinetic Modeling

To obtain a meaningful kinetic representation, the timescales can be extracted using a model based on knowledge of the system. We applied global target analysis (GTA) that consists of compartments, each representing a kinetically distinguishable state with spectral features, known as species associated difference spectra (SADS), with an associated lifetime. The number of compartments (lifetimes) as well as their temporal ranges were based on the lifetime distributions obtained by LDA (see **figure s16** and **table s4**). Because our focus lies on the characteristic differences between the variants, we applied the same model for all samples. Specific lifetime values were obtained by exponential fitting of the ΔΔ datasets (**table s5**) and time-resolved fluorescence data and were kept soft-fixed (with a 5-10% margin) within the GTA. The best fit was achieved by a five-compartment model (**figure 4.C, figure s14.C** and **table s6**) with equilibria between compartments S1 and S2, as well as S2 and S3. The SADS of S1 mainly reflect the unequilibrated bulk antenna with strong ESA features below and above the GSB of the core antenna. These ESA signatures are less pronounced in S2 due to the progressed energy equilibration, which also leads to a broadening and a slight redshift of the GSB due to the contributions by red and P700*. The transition from S2 to S3 occurs with τ_2_ (WT < RedB < RedAB < RedA) and is reflective of two processes: the population of red and the initial formation of the charge separated state. The differences in lifetimes show a significant site-dependent effect on the kinetics at this stage. The prolonged lifetimes and the strong absorption bands around 705-715 nm in RedA and RedAB (**figure 4.C** and **figure S14.C**) also underline an accumulation of the red absorbing species, which was already suggested by the ΔΔ spectra. Charge separation is indicated by the distinct absorption feature at >725 nm and the superimposed (with the GSB) absorption band at ∼695 nm, which can be attributed to the formation of the neutral Chl_690_ (Savikhin et al., 1999, 2001; Schaffernicht C Junge, 1982). As inferred from the LDA (**figure s16** and **s17**), these processes are temporally and spectrally too close to separate into individual compartments. The S4-SADS reflect the progressed charge separation which is characterized by further accumulation of the Chl_690_ absorption at ∼695 nm. Interestingly, this transition is facilitated in RedB (7.2 ps) compared to the WT (8.5 ps) or the variants RedA (9 ps) and RedAB (12 ps) (τ_3_, **table s6**), again showing a site dependent effect, this time probably related to charge separation processes. The final spectrum (S5) represents the completed charge separation with the characteristic features of the P700+/A_1-_ species (including the blue-shifted and fully accumulated Chl_690_ band) and is formed within 30-38 ps.

We found two distinct effects associated with different red site insertions in PSI. RedA, which is positioned far from P700, leads to strong, red-shifted GSB with a maximum amplitude around 10 ps (**figure 4.A** and **B, solid lines**), which probably represents localization of excitation to the RedA chls (mixed with other red sites). In contrast, RedB, which is positioned near to P700, leads to a rapid (<10 ps) decrease in the antenna GSB (**figure 4.A** and **B, dashed lines**), which probably represents rapid energy localization to the RedB chls followed by fast transfer from RedB to the PSI ETC. The double difference spectra kinetic analysis suggests that the double mutant contains both the faster process induced by RedB (τ_2_, **table s5**) and the slower processes induced by RedA (τ_3_, **table s5**).

**Figure 5.**
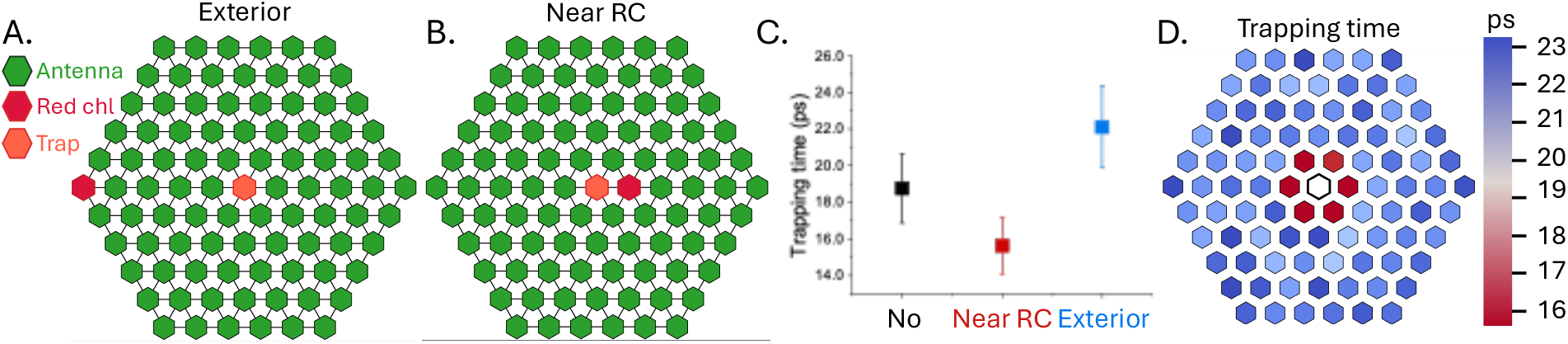
The effect of red site location on trapping time in an idealized antennae system. **A**. and **B**. Two different scenarios for red chl location were tested on a hexagonal array. An exterior or a RC proximal red chl location (‘A’ and ‘B’ respectively. **C**. Simulating the trapping rate for both red site location scenarios at an energy value of 600 cm^-1^ (the estimated energetic gap for both the RedA and RedB mutants) shows significant differences compared to a no energy gap scenario, with the near RC scenario leading to significantly shorter and the Exterior scenario leading to significantly longer trapping times. **D**. Repeating the scan for all possible well locations using an energetic bias of 600 cm^-1^ shows that on the examined lattice, positional effects have sharp boundaries.

## Discussion

In this work, we explored the relationship between the location of low energy sites and their effect on primary processes in the core antenna of PSI. Typically, the red chl in PSI are bundled together and treated as one or two pools because a direct assignment of transient spectral changes to a specific red chl site is not known (Gobets, Kennis, et al., 2001; Gobets, Van Stokkum, et al., 2001; Gobets & van Grondelle, 2001; Kaucikas et al., 2017; Molotokaite et al., 2017; Slavov et al., 2009). In our study, however, we can make such an assignment by direct comparison between the mutants to isolate the red chl contribution from a specific site.

Our time-resolved data suggest that different red site locations lead to significant differences between the ultrafast kinetics of the core PSI antenna. There are clear differences between RedA and RedB despite both generating similar energetic shifts in their respective ground state spectrum as inferred from the absorption difference spectra (**figure 4**, far right, solid gray line**, figure s15**). The effects are best visualized in the ΔΔ spectra (**figure 4.A, right panel**, which reduces the transient spectral changes to the contributions solely stemming from the mutations. The RedA and RedB sites are connected to the RC via the bulk antenna network and therefore take part in the later phases of the energy equilibration (compartments S2 and S3 in our model, **figure 4.C**). Direct transfer from RedA to any of the RC chl is highly unlikely given the large distance separating these entities (∼48 Å, **figure 1.A**). However, in case of RedB, only one or two transfer steps separate the PSI ETC from RedB given the small distance (∼29 Å, **figure 1.A**). The presence of red chl at site A (in RedA and RedAB) slows down energy transfer to the RC (τ_3_, **table s6**) and consequently leads to a delayed trapping time (τ_4_, **table s6**) compared to the wildtype and RedB. The presence of red chl at site B, on the other hand, speeds up energy transfer to the RC (τ_3_, **table s6**) and shows a smaller effect on trapping time (τ_4_, **table s6**). These effects suggest that red traps closer to the RC/ETC enhance the efficiency of the core antenna whereas red traps further from the RC/ETC significantly delay trapping.

We simulated these two scenarios in an idealized photosynthetic antenna model arranged as a set of 90 nodes on a hexagonal array. Similar models have been useful in evaluating the relationship between system parameters, such as the dependency of the trapping time on the number of nodes (Beauregard et al., 1991; Pearlstein, 1982).

Simulating trapping in idealized systems containing “RC proximal” or “RC distal” red chls showed the same trend observed in our measurements. These results indicate that the observed trends do not depend on the precise location of the red sites but reflect a general property of antennae. Similarly, these results are robust to a range of energy gaps in simulations (**figure s19**).

The energetic configuration of photosynthetic systems is sometimes easily rationalized, for example the cyanobacterial phycobilisome is a classic example of energetic funneling or the FMO protein has been called an energetic ‘wire’ (Adolphs & Renger, 2006; Glazer, 1985). In case of the PSI core, which is a dense, highly coupled chl array, the location of low energy or red chls is more challenging to rationalize a priori (Croce & Van Amerongen, 2013). There are some indications that red sites in the PSI supercomplexes are located at the interfaces of LHC antennae and the PSI core in higher plants (Ben-Shem et al., 2003; Croce et al., 2002).

The location of RedA in the cyanobacteria PSI, which is a naturally occurring variant, initially appears counterintuitive as it is not clear how it contributes to PSI function. We suggest that in the context of larger arrays, such as PSI-IsiA, RedA can function similarly to RedB, i.e, as a RC proximal red site in the context of a much larger PSI-antennae array. It should also be noted that although idealized simulations show a decrease in trapping time in the RC proximal scenario, this is not what we observed in the RedB mutation, which still impacts the trapping time in PSI. It is possible that in the context of PSI, RedB represents the best balance between increasing light harvesting into the far-red region while minimizing the impact on trapping kinetics. It is also possible that in their native context, both RedA and RedB are accompanied by further energetic modifications in the core antenna which decreases their impact on trapping time. Finally, it is possible that trapping time is not the best indicator of PSI “efficiency” (for lack of a better term) and a different metric, for example minimizing the productions of chl^3^, determines the “cost” of red sites and their location.

Our analysis of the RedB site across a series of PSI structures suggests that the 6803 RedB site can adopt a red configuration. This information can only be observed when there are multiple, high resolution, structures of the same region in PSI. At lower resolution, differences of fractions of an Å, which can significantly change the electronic properties of chls, cannot be determined with high confidence (Lin et al., 2024). Going back to room temperature and the native context of PSI, it is hard to evaluate the time the wt 6803 RedB site spends in this configuration, but given the spectroscopical evidence, the majority of 6803 PSI trimers are not in this configuration. An important question is whether the RedB site is unique in its ability to adopt multiple conformations or does this property exist in a few or all chl is PSI. Our results suggest, but do not prove, that other chl dimers may be heterogenous. Single complex observations of PSI suggest a considerable variability in its emission spectra (Jelezko et al., 2000; Riley et al., 2007), which fits well with red sites heterogeneity. The current picture is undoubtedly partial with regards to conformational variability of chls in PSI, but It suggests that the best way to discuss red chl in PSI is as a distributed property that can be associated with different sites to different extent.

In conclusion, there is an inherit balance between the benefit of an energetic gap contained in an antenna system and the impact of such a gap on the performance of the system. Here, we examined one aspect of this balance: the location of red site in the antenna of PSI and its impact on the photophysics of the system. We find that the deleterious impact of an energetic well on the performance of the system are minimized by placing this well in an RC proximal position. In the context of cyanobacteria, we suggest that placing the well in an RC distal position can be rationalized in the PSI-IsiA arrays that are common in nature. In such systems, these sites can be considered as ‘PSI-core proximal’ and we expect that their impact on trapping time would be similarly minimized.

## Acknowledgements

This work was supported by the U.S. Department of Energy, Office of Science, Office of Basic Energy Sciences, Division of Chemical Sciences, Geosciences, and Biosciences under Award \#DE-SC0018097 (to G.S.S.-C) and by the Department of Energy, Office of Science, Office of Basic Energy Sciences, Division of Chemical Sciences, Geosciences, and Biosciences under Award \#DE-SC0022956 (to Y.M). M.A also acknowledges the German Research Foundation (DFG) for their generous financial support under the Walter Benjamin Programme (award number 564394927). MR acknowledges support from the U.S. Department of Energy, Office of Science, Office of Basic Energy Sciences, Division of Chemical Sciences, Geosciences, and Biosciences under Award \#DE-SC0022884. The cryo-EM work utilized a Selectris X imaging filter and a Falcon 4i direct electron detector (DED) that was purchased through funding from a National Institutes of Health Shared Instrumentation Grant (SIG; 1S10OD036204-01).

## Data availability

The cryo-EM density maps generated in this study have been deposited in the Electron Microscopy Data Bank (EMDB) under accession code: PSI monomer masked map-EMD-71005 [https://www.ebi.ac.uk/emdb/EMD-71005], PSI trimer refined in C1 map-EMD-71048 [https://www.ebi.ac.uk/emdb/EMD-71048]. The atomic coordinates generated in this study have been deposited in the Protein Data Bank (PDB) under the accession codes: PSI monomer - 9OY9 [https://doi.org/10.2210/pdb9OY9/pdb], PSI trimer in C1 - 9OZ3 [https://doi.org/10.2210/pdb9OZ3/pdb]. Source data are provided with this paper.

## Notes

### Competing Interest Statement

The authors have declared no competing interest.

