## supplementary data for "Red states modulate energy localization between the antenna and trap in PSI"

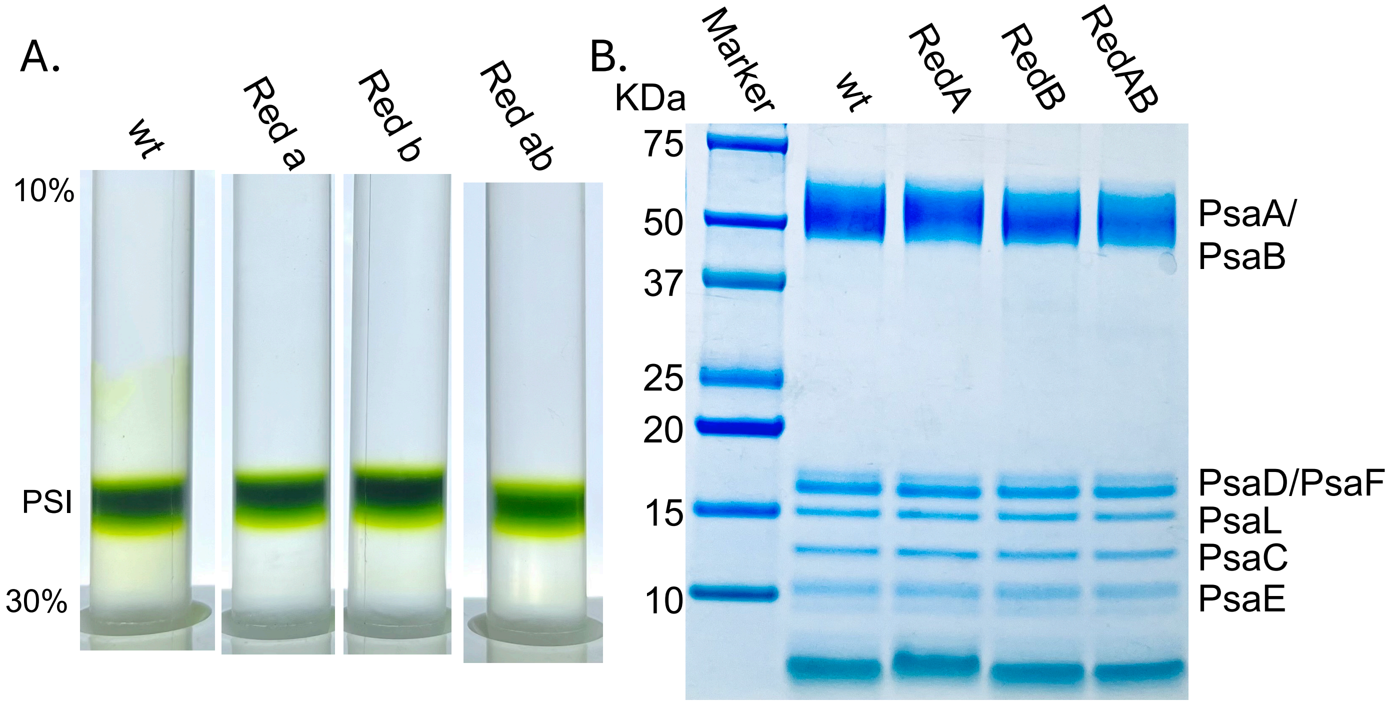

**Figure S1: Purification of PSI complexes**. **A**. The isolated PSI complexes from wt, RedA, RedB, and RedAB were collected from the chl containing band of the 2$^{nd}$ sucrose gradient. **B**. SDS-PAGE gel of the purified PSI from wt, RedA, RedB and RedAB that were collected

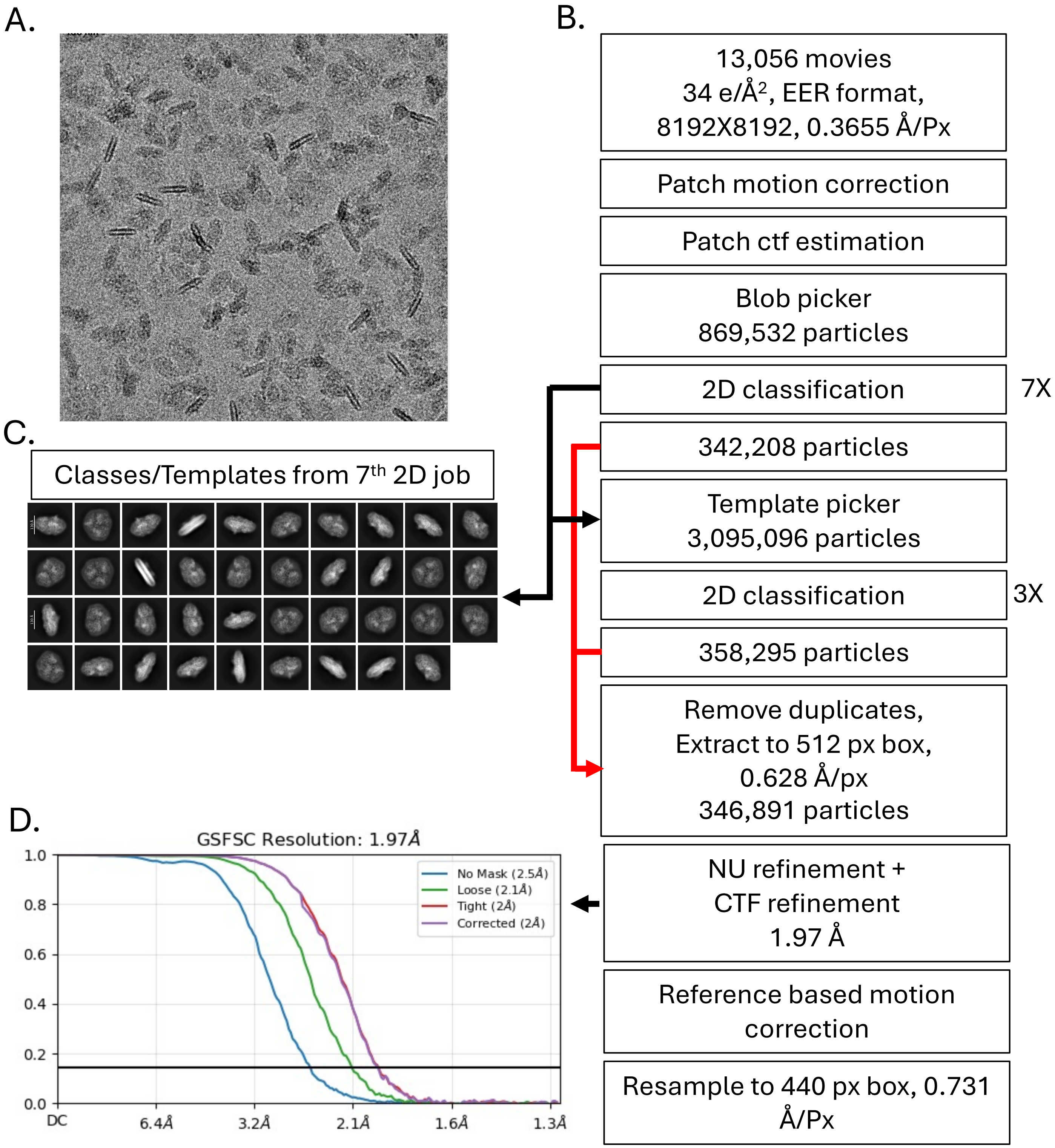

**Figure s2:** CryoEM data processing. **A**. Sample micrograph after motion correction. **B**. different stages of data process, all carried out using CryoSPARC. C. 2D classes after the initial screening for "bad" particles that were used for a 2nd round of template picking. **D**. FSC curves before referenced based motion correction.

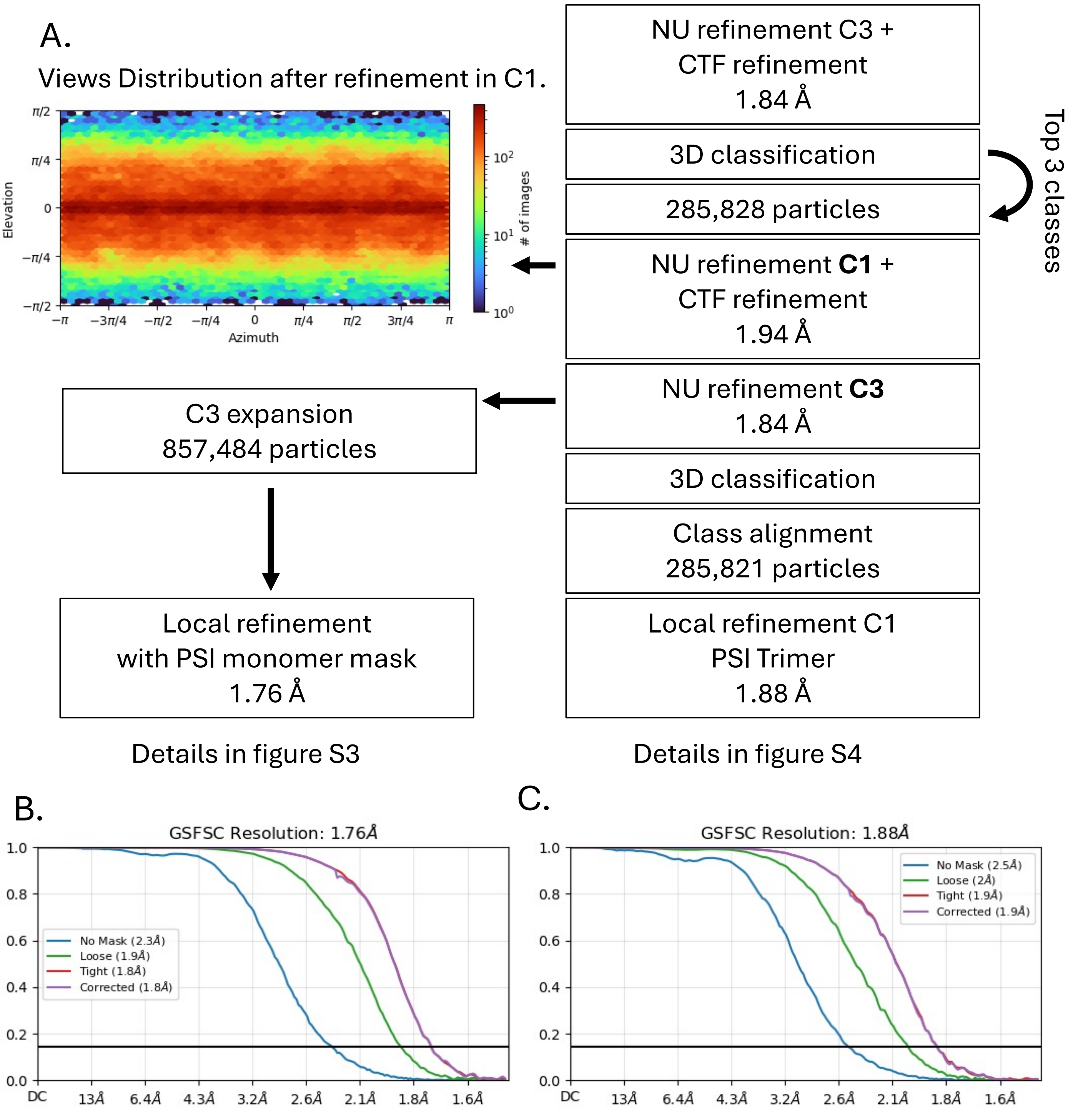

**Figure s3**: CryoEM data processing – final volumes used for model building. **A**. Processing steps to obtain the final particles dataset and the maps used in model building, all done in CryoSPARC . **B**. FSC for locally refined monomer after C3 expansion. **C**. FSC for trimer refined in C1.

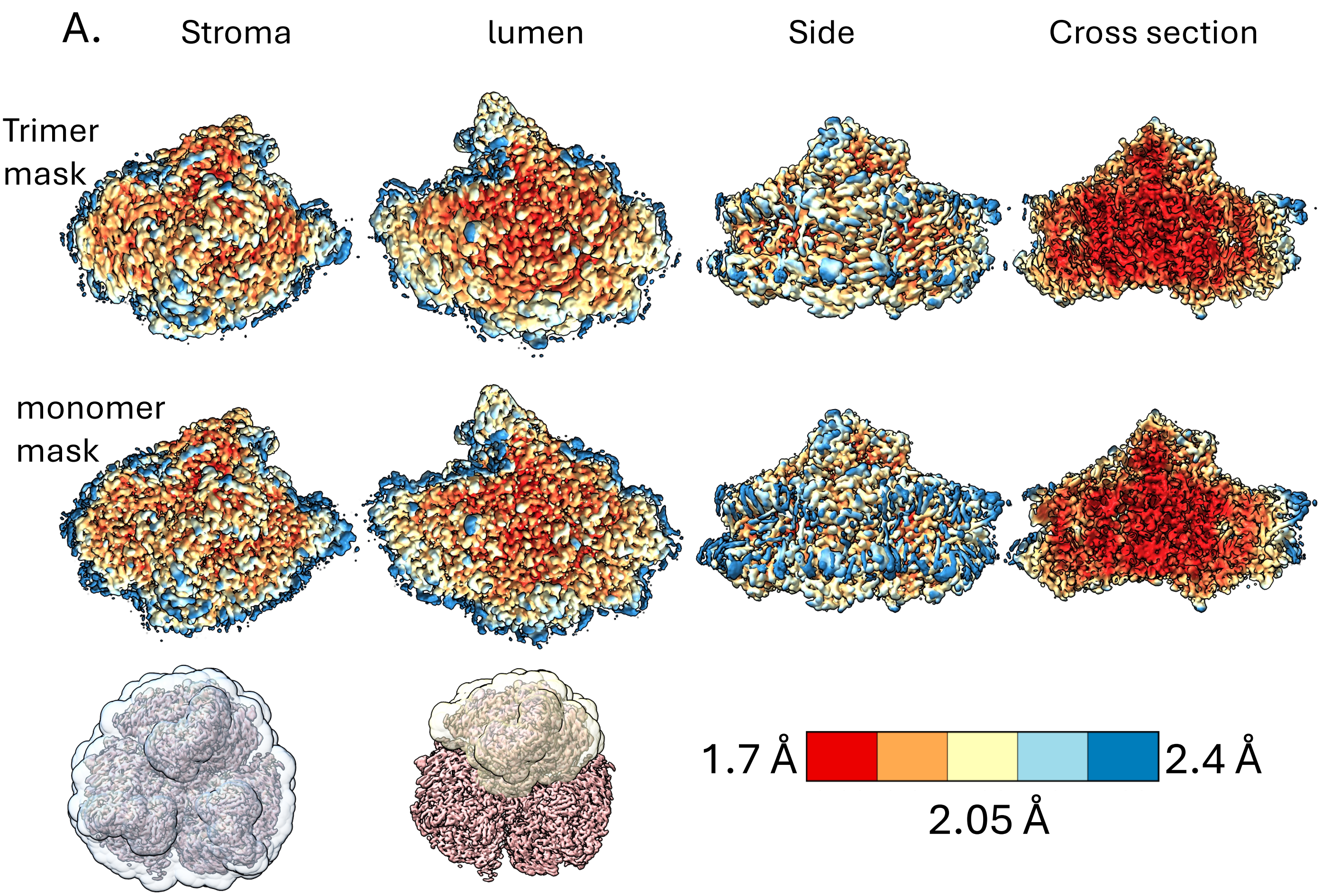

PDB ID 9OY9, Extended PDB ID pdb_00009OY9, EMD-71005
10.2210/pdb9oy9/pdb

D_1000296570

**Figure s4**: Local resolution of the C3 expanded dataset, before (top row) and after (bottom row) focused refinement. The masks used in refinement are shown at the bottom row, next to the resolution scale bar. Monomer model and map deposited as PDB ID 9OY9, EMD-71005.

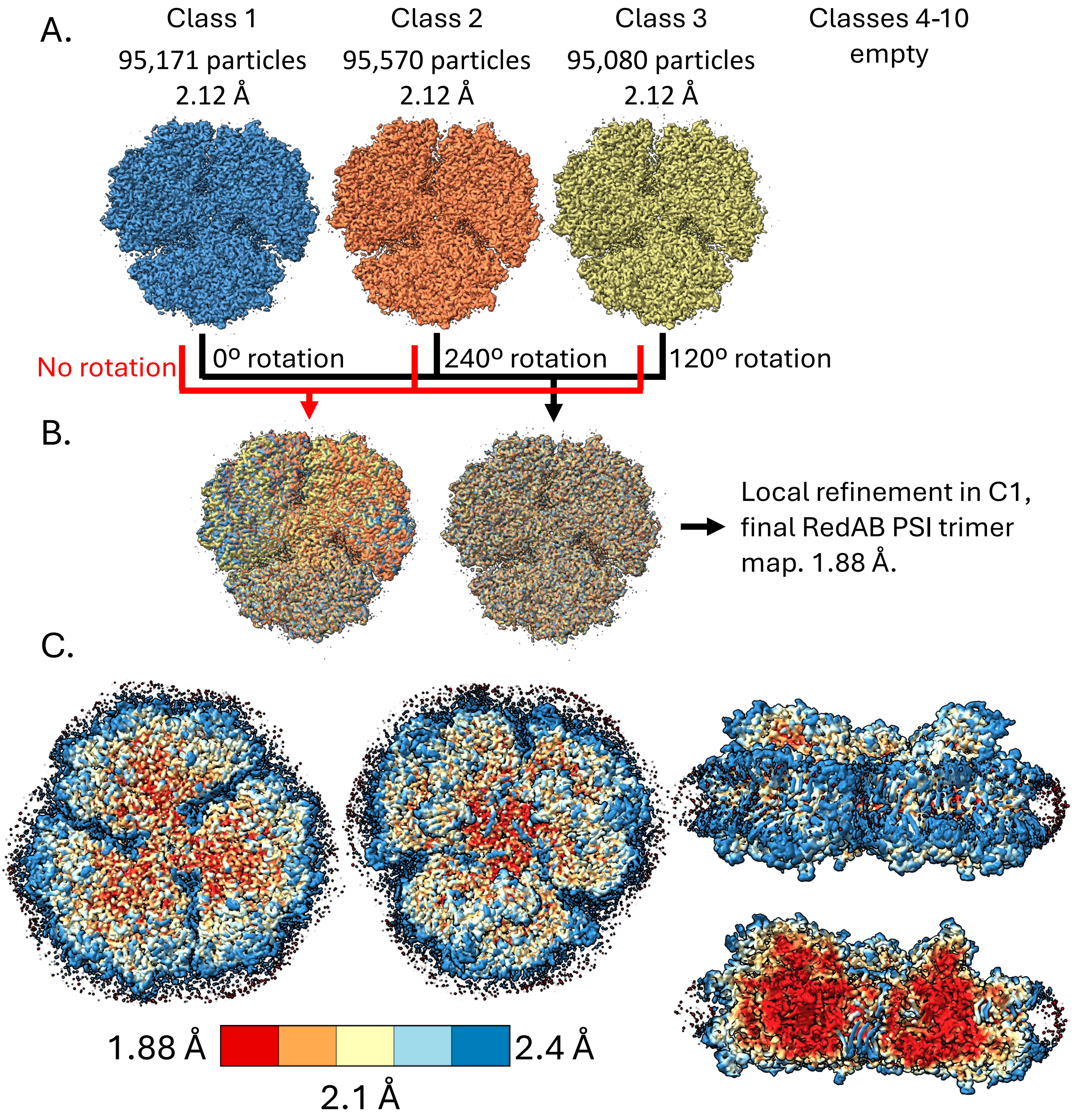

**Figure s5:** Classification of different trimer conformations. A. Three main classes at equal proportions were obtained after 3D classification without alignment of the entire dataset. Without rotations the three classes could not be superposed (red arrows, B), but with 120º or 240º rotations, the classes were superposed (black arrows, B). These alignments were then applied to the particles in each class resulting ~0.1Å improvements in the overall resolution of the C1 trimer map and the elimination of much of the heterogeneity observed at the periphery of the PSI trimer, as seen in the local resolution map C. C1 trimer deposited as PDB ID 9OZ3, EMD-71048.

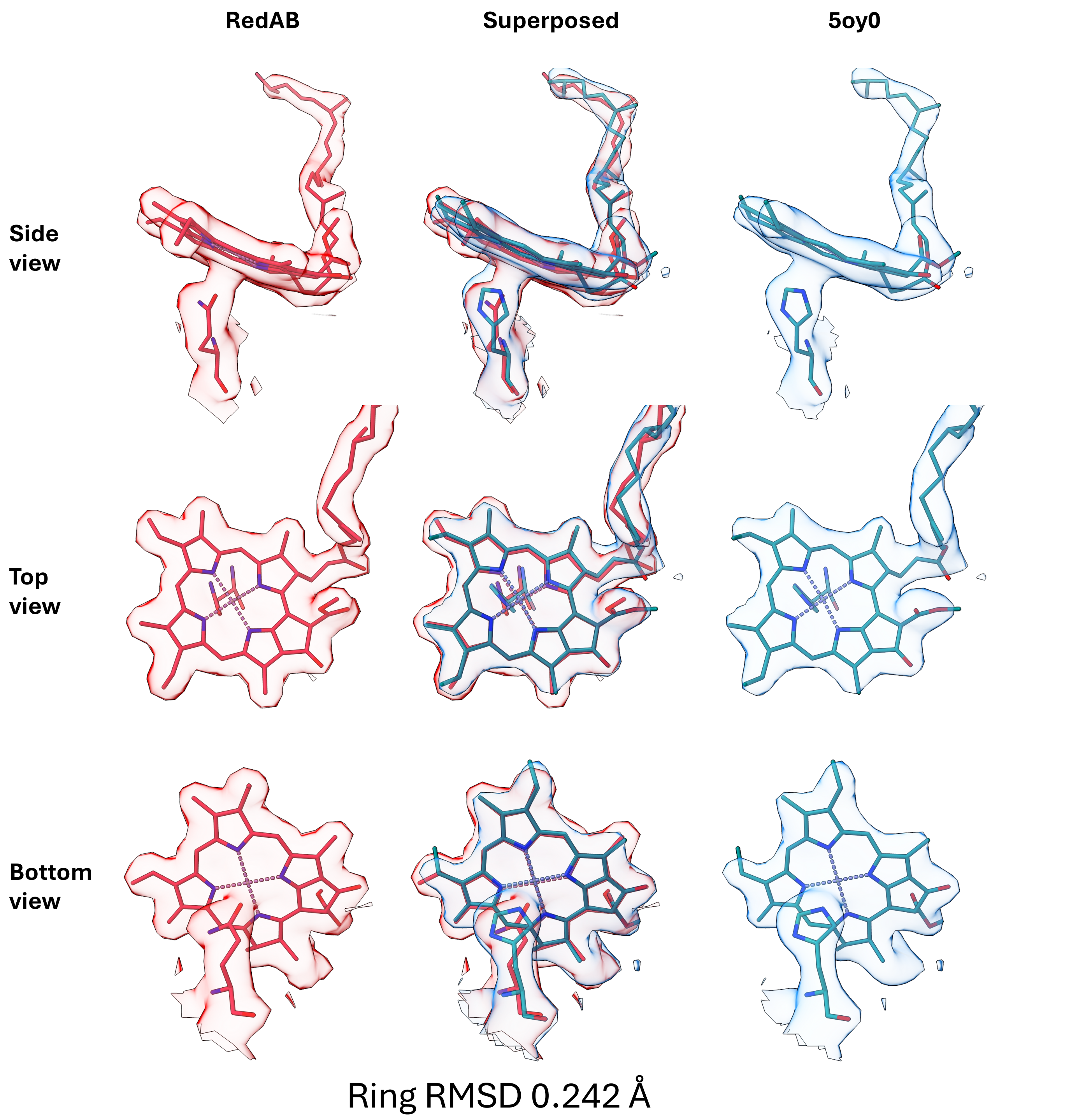

**Figure s6**: Chlorophyl B7 map and model superposition between RedAB and the 6803 wt crystal structure (5oy0) obtained at 2.5 Å resolution. The RedAB map and model in red, 5oy0 map in blue and model in teal. Map superposition was done in Chimerax on the RedAB cryoEM map. Models were superposed using PsaB and the matchmaker command in Chimerax. Maps are drawn around the mutated amino acid and the entire B7 chl. For scale, bond thickness in these images is set to 0.1 Å. Map differences around the mutated side chains are clearly seen. The Oxygen atom of the mutated Gln residue occupies nearly the same position as the original Nitrogen atom in the His imidazole group (separated by 0.22 Å) and the two chls are nearly perfectly superposed with an RMSD of 0.242 Å measured across all chl ring atoms.

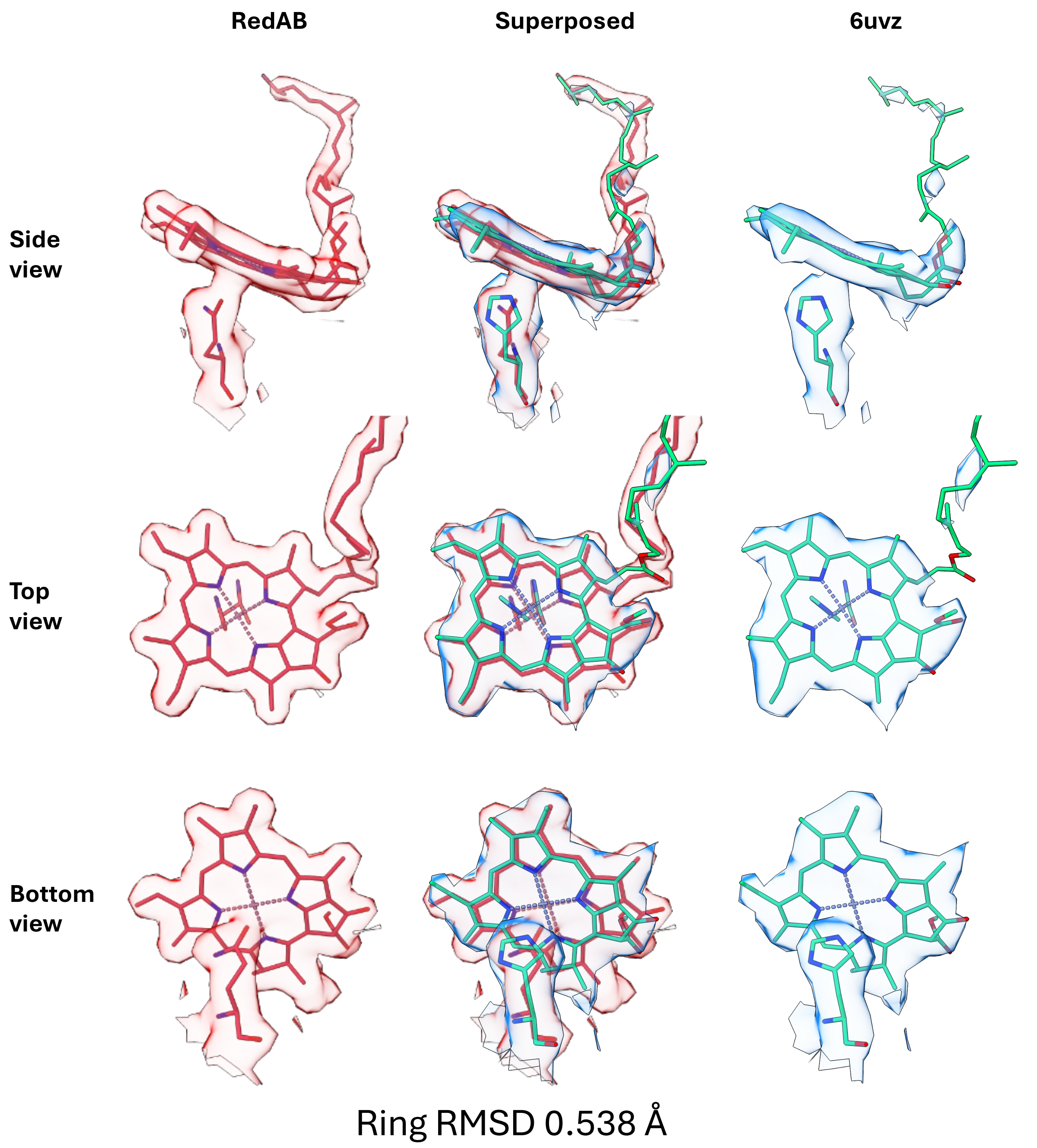

**Figure s7**: Chlorophyl B7 map and model superposition between RedAB and the RedA CryoEM structure (6uvz) obtained at 3.1 Å resolution). The RedAB map and model in red, RedA map in blue and model in light green. Map superposition was done in Chimerax on the RedAB cryoEM map. Models were superposed using PsaB and the matchmaker command in Chimerax. Maps are drawn around the mutated amino acid and the entire B7 chl. For scale, bond thickness in these images is set to 0.1 Å. The lower resolution of the RedA map is appearent, the distance between the nitrogen atoms of the Gln and His residues is larger (0.67 Å) and a larger chl shift is clearly observed (ring RMSD of 0.538 Å).

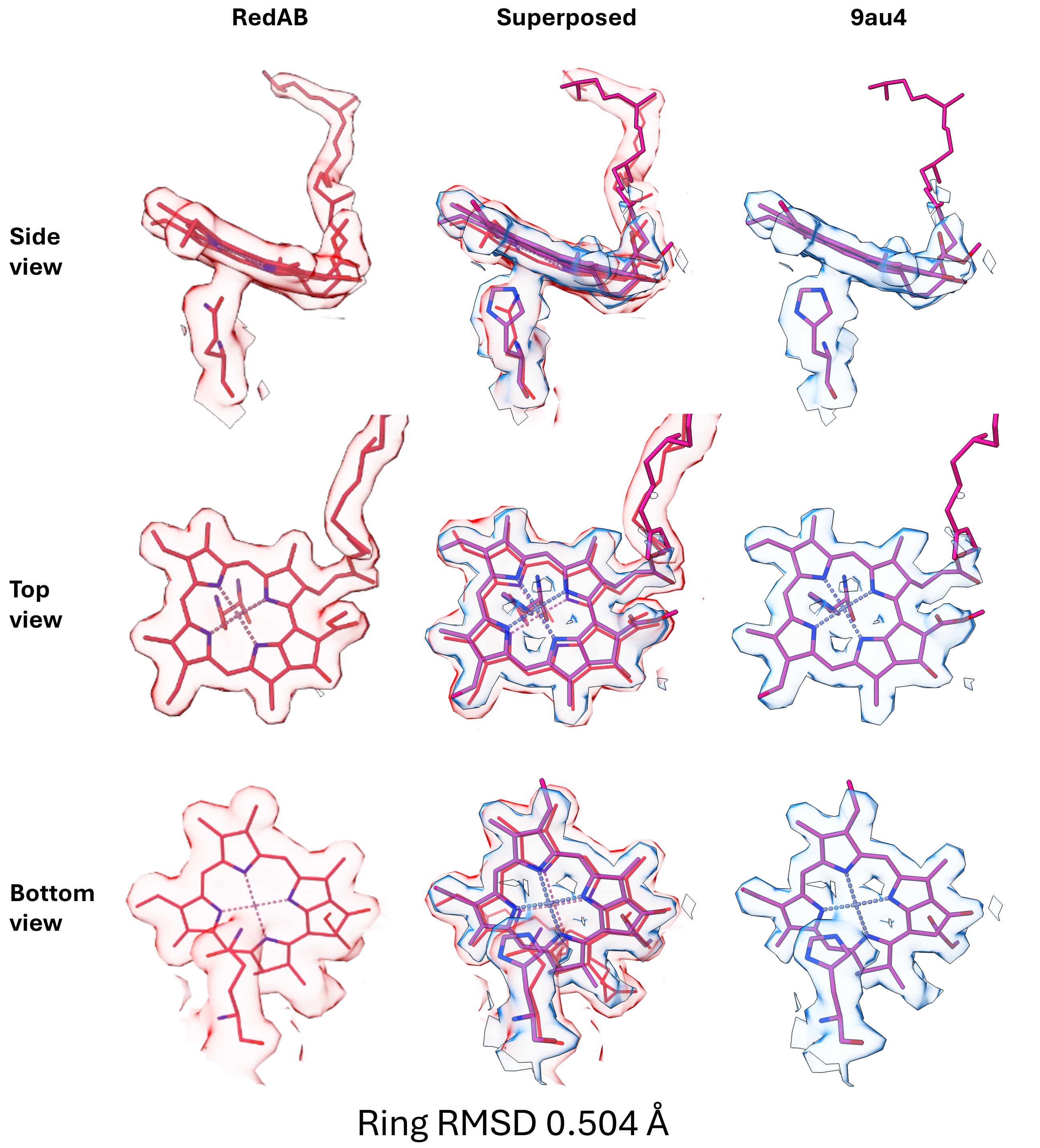

**Figure s8**: Chlorophyl B7 map and model superposition between RedAB and another reference 6803 CryoEM structure (9au4) obtained at 2 Å resolution). The RedAB map and model in red, 9au4 model in grey. The models were superposed using PsaB and the matchmaker command in Chimerax. Similar shifts to those observed in the RedA case are seen, with a total chl ring RMSD of 0.504 Å. The map is drawn around the mutated amino acid and the entire B7 chl. For scale, bond thickness in these images is set to 0.1 Å.

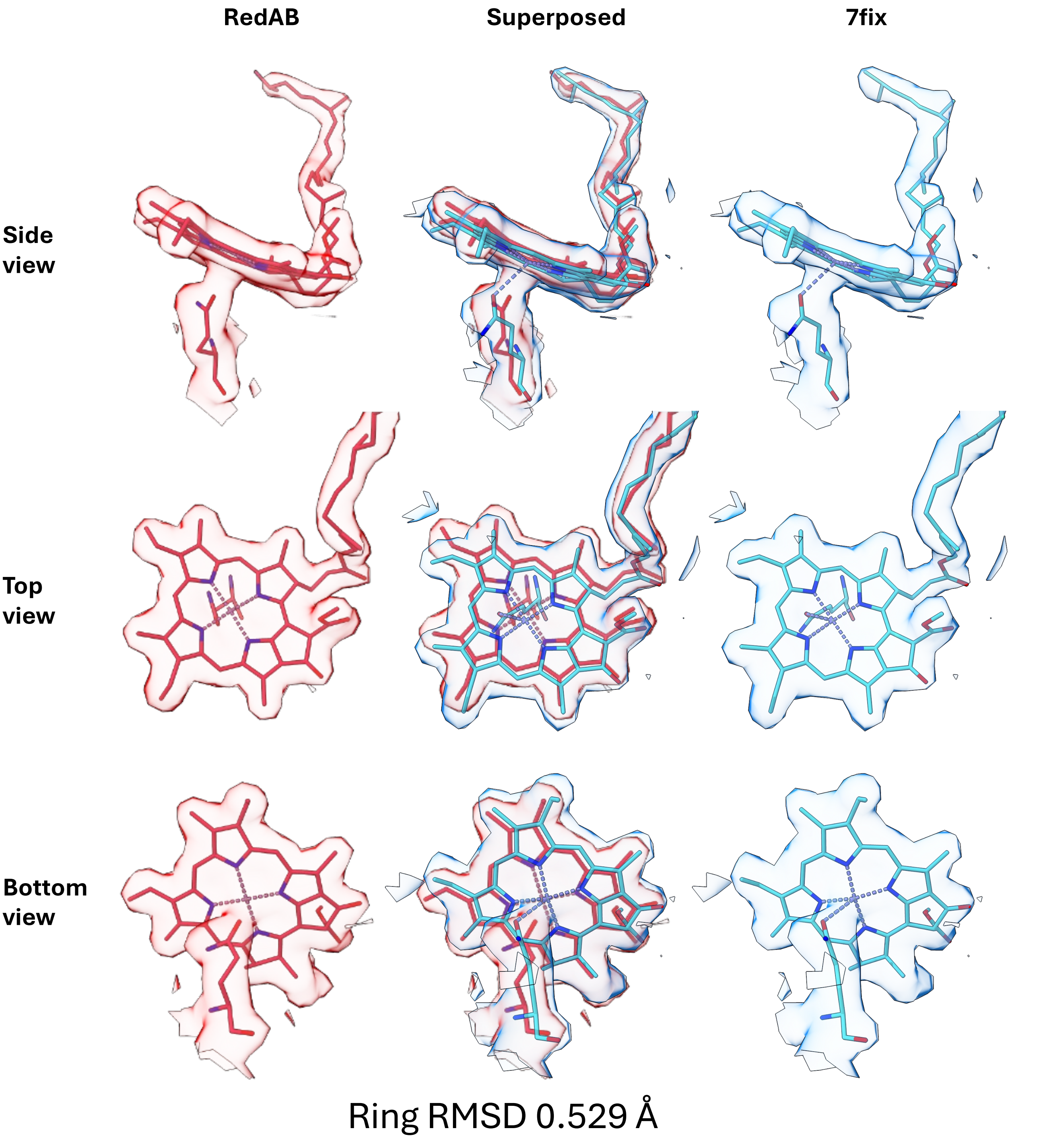

**Figure s9**: Chlorophyl B7 map and model superposition between RedAB and the TV CryoEM structure (7fix) obtained at 1.97 Å resolution). The RedAB map and model in red, TV map in blue and model in cyan. Map superposition was done in Chimerax on the RedAB cryoEM map. Models were superposed using PsaB and the matchmaker command in Chimerax. Maps are drawn around the mutated amino acid and the entire B7 chl. For scale, bond thickness in these images is set to 0.1 Å. Maps are of comparable resolution, the coordination for chl B7 was modeled differently in 7fix making distance measurements between the Nitrogen atoms irrelevant. The clear shift in chl position is in the opposite direction to the one observed in RedA with a ring RMSD of 0.529 Å.

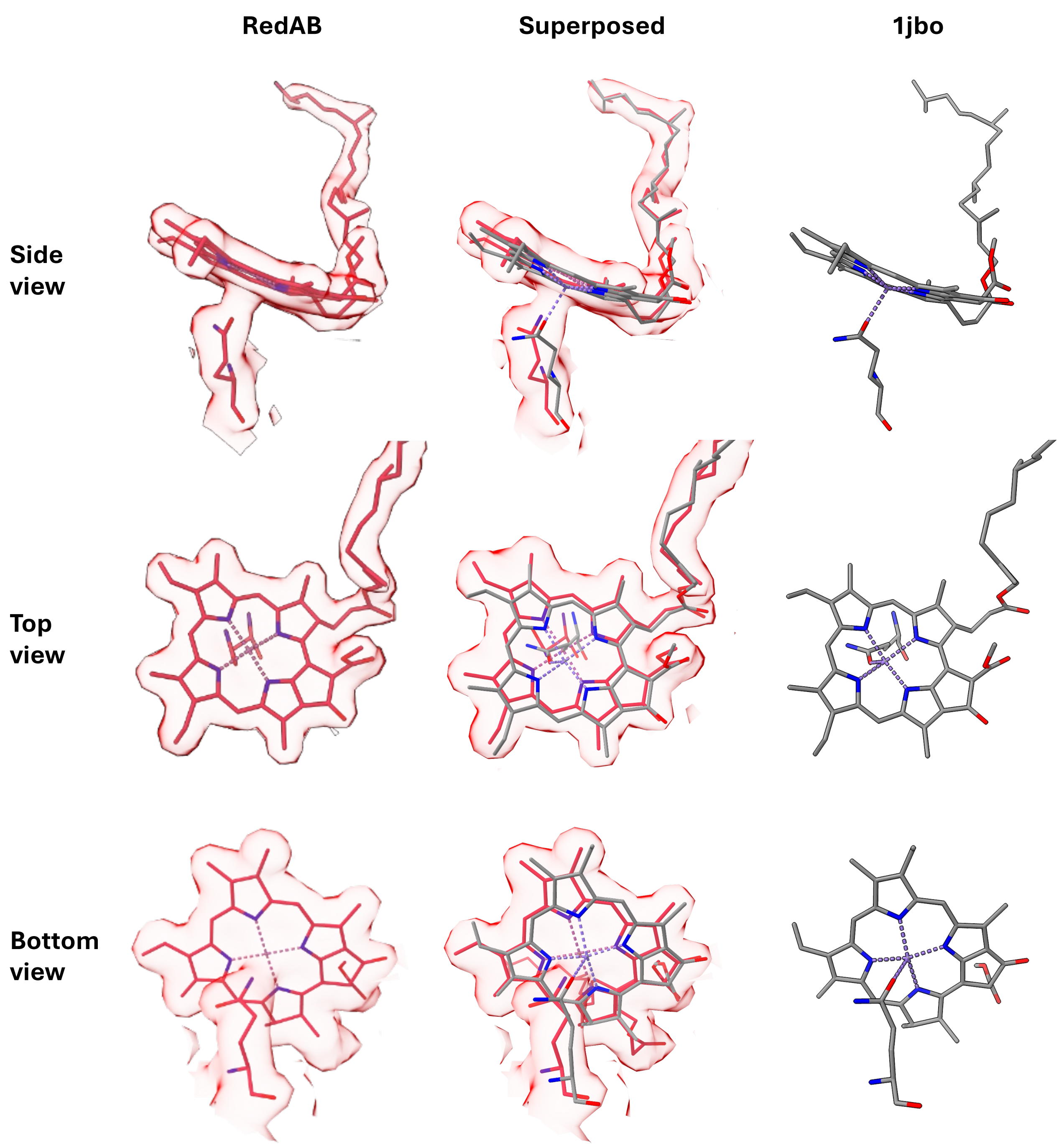

Ring RMSD 0.495 Å

**Figure s10**: Chlorophyl B7 map and model superposition between RedAB and the TV crystal structure (1jb0) obtained at 2.5 Å resolution). The RedAB map and model in red, TV model in grey (no publicly available map). The models were superposed using PsaB and the matchmaker command in Chimerax. The map is drawn around the mutated amino acid and the entire B7 chl. For scale, bond thickness in these images is set to 0.1 Å. Onc clear difference is the conformation of the axial Gln residue, in 1jb0 the oxygen is used as the coordinating atom. The two possibilities are difficult to distinguish, in the RedAB model the Nitrogen atom was used as the axial ligand based on better map fit, the distance between the coordinating atoms is 0.46 Å. The position of B7 is clearly closer to 7fix with a chl ring RMSD of 0.495 Å between RedAB and 1jb0.

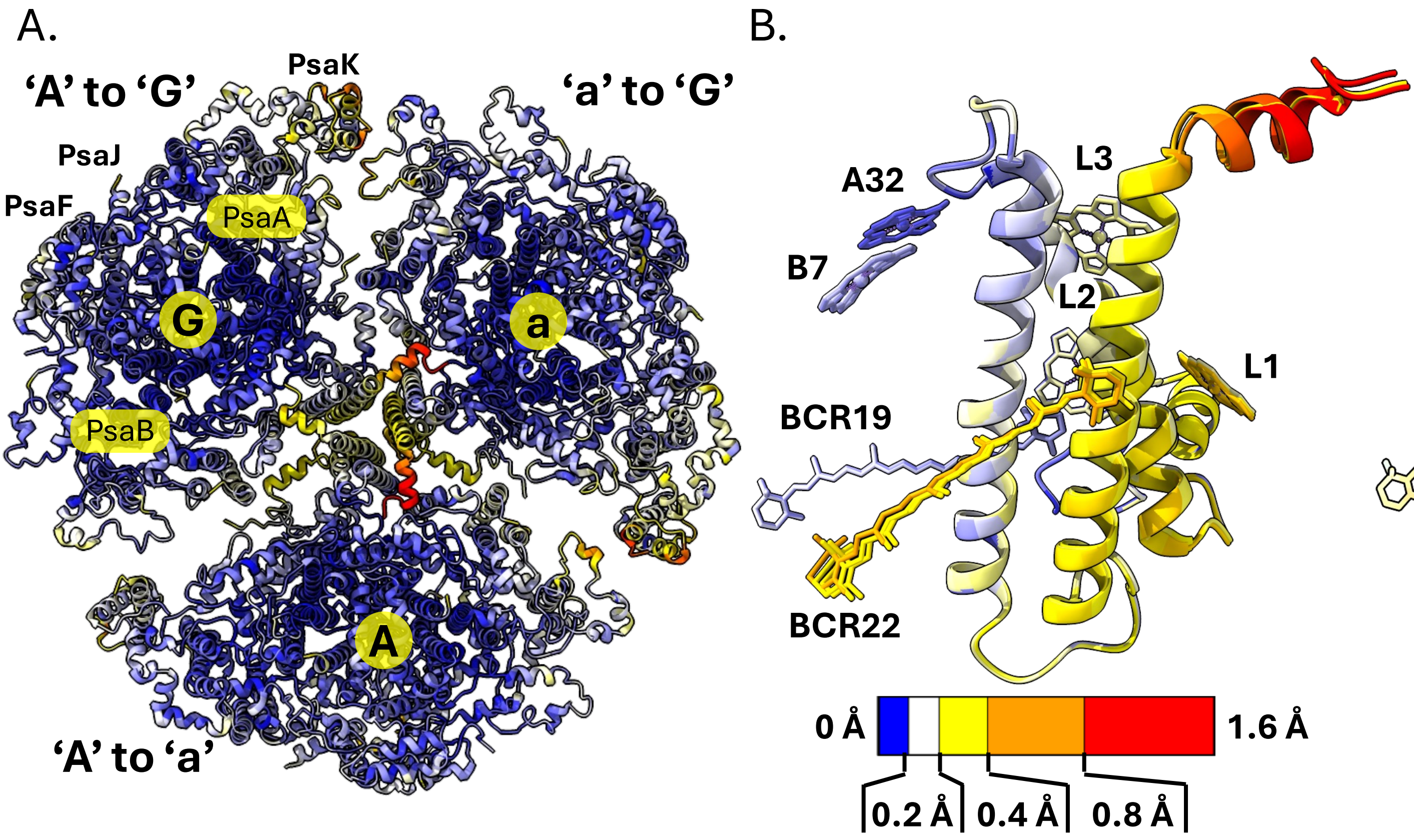

**Figure S11. Structural heterogeneity across the PSI trimer. A.** Cartoon representation of the PSI trimer colored based on RMSD between the different monomers. Monomers are identified based on their PsaA chain ID ('A', 'a' or 'G'). Each monomer is colored based on the RMSD (or simply the distances) between the C alpha atoms across all residues. The pair is identified in the figure, for example, monomer G is colored based on the RMSD between monomer 'A' to 'G'. this identified monomers 'G' and 'A' as being similar to each other, with monomer 'a' being somewhat different, the most noticeable differences are a slight shift in the orientation of the PsaL C terminus. **B**. PsaL and the RedB site chl's colored according to their RMSD, all three PsaL subunits were superposed to visualize the differences among them. **B**. Same as 'B' only colored based on local resolution.

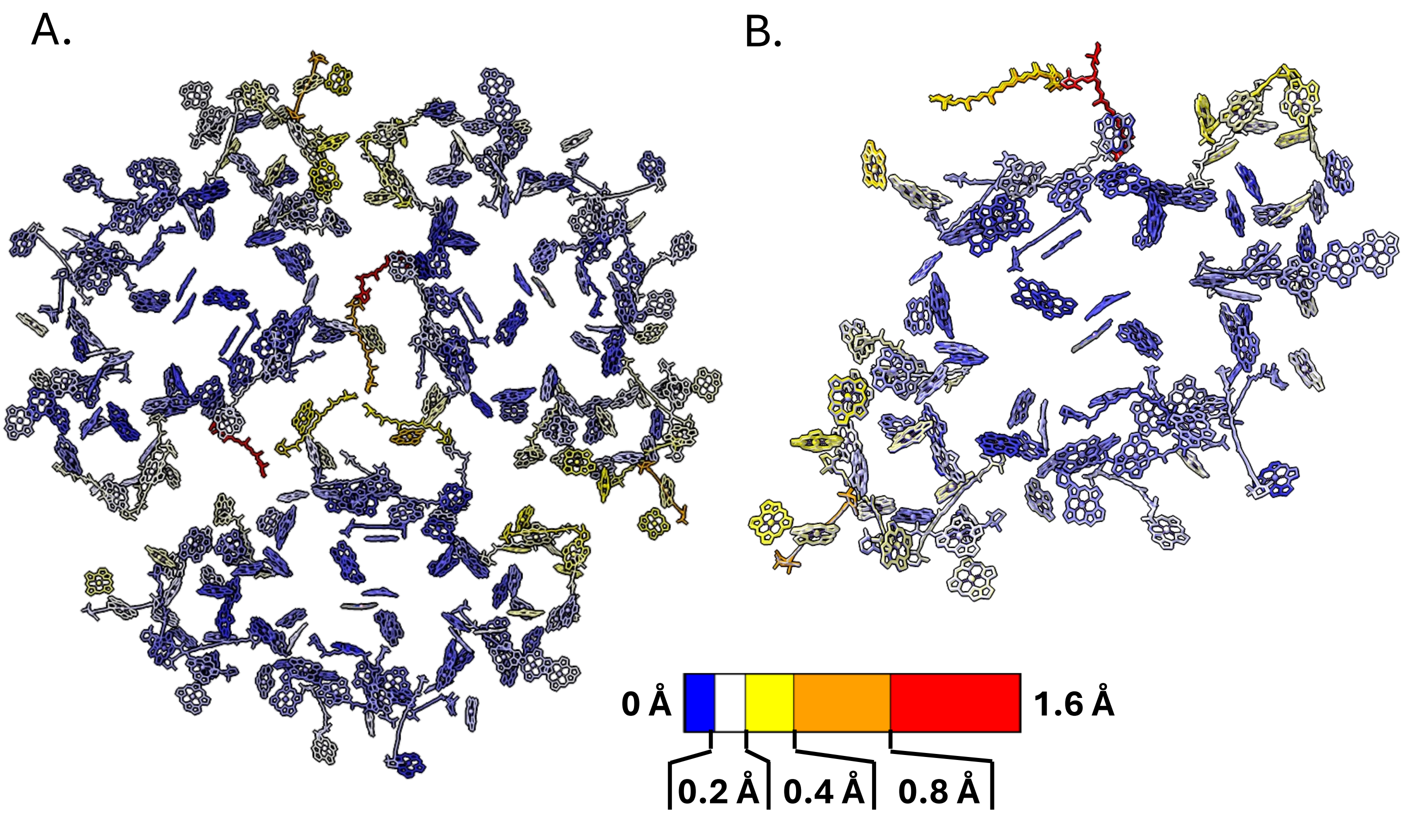

**Figure s12 :**A. PSI monomers colored by rmsd (calculated over CA only). Showing that PsaL in the ‘a’ monomer adopts a somewhat different conformation. In addition, PsaK, PsaF and PsaA show higher rmsd values compared to the mean of the entire complex. B.

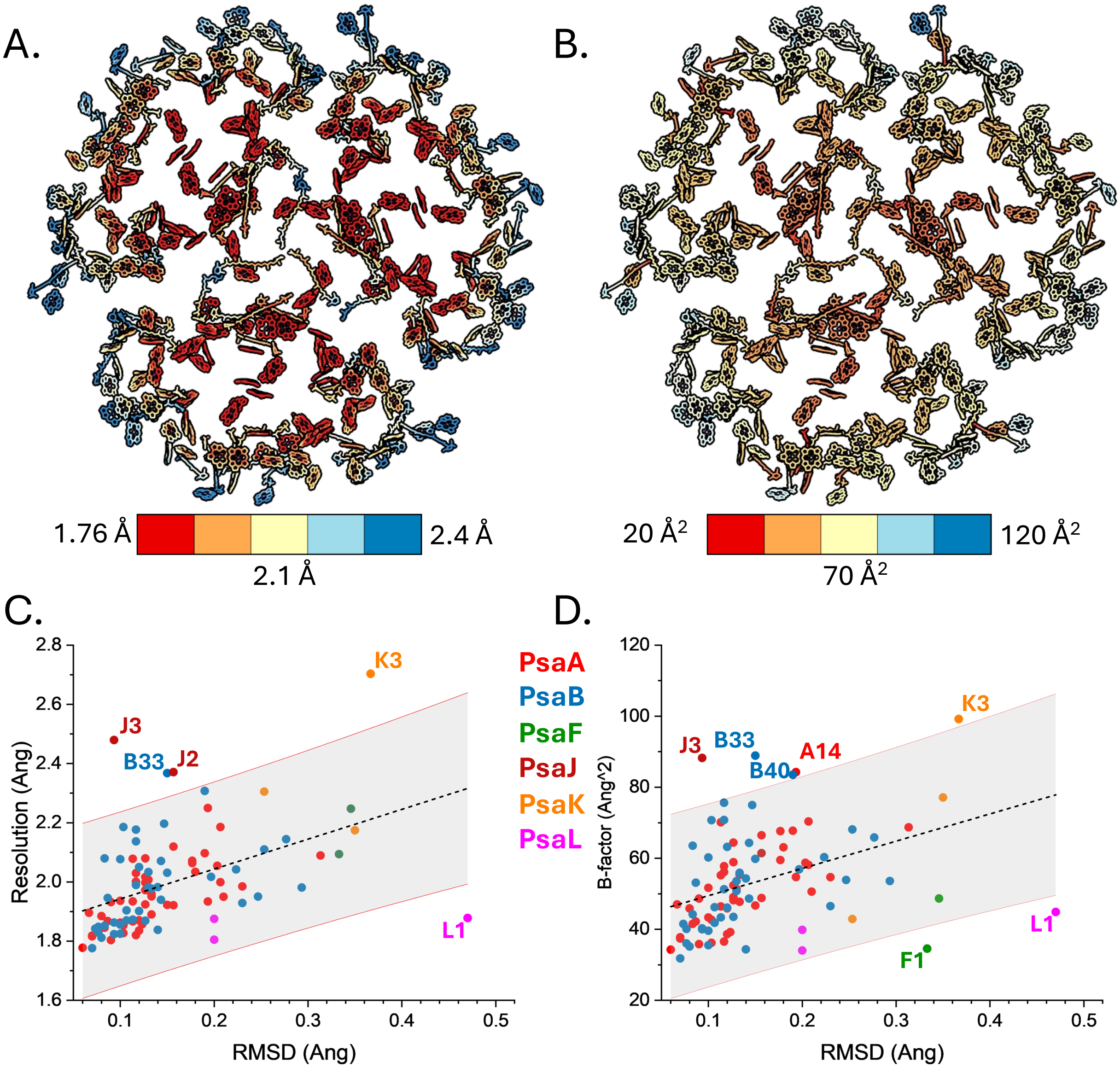

**Figure s13**: **Positional uncertainty and structural variability across the PSI chls**. **A**. PSI chls and carotenes colored according to their local map resolution. **B**. PSI chls and carotenes colored according to their average b factor for the ring atoms. **C**. RMSD (positional variability) plotted against local resolution (chls averaged across trimers for both measures). Pearson's correlation coefficient is 0.44. **D**. RMSD (positional variability) plotted against b factor (chls averaged across trimers for both measures). ). Pearson's correlation coefficient is 0.39. In both cases most chlA RMSD values fall within 95% confidence intervals and may be explained by resolution.

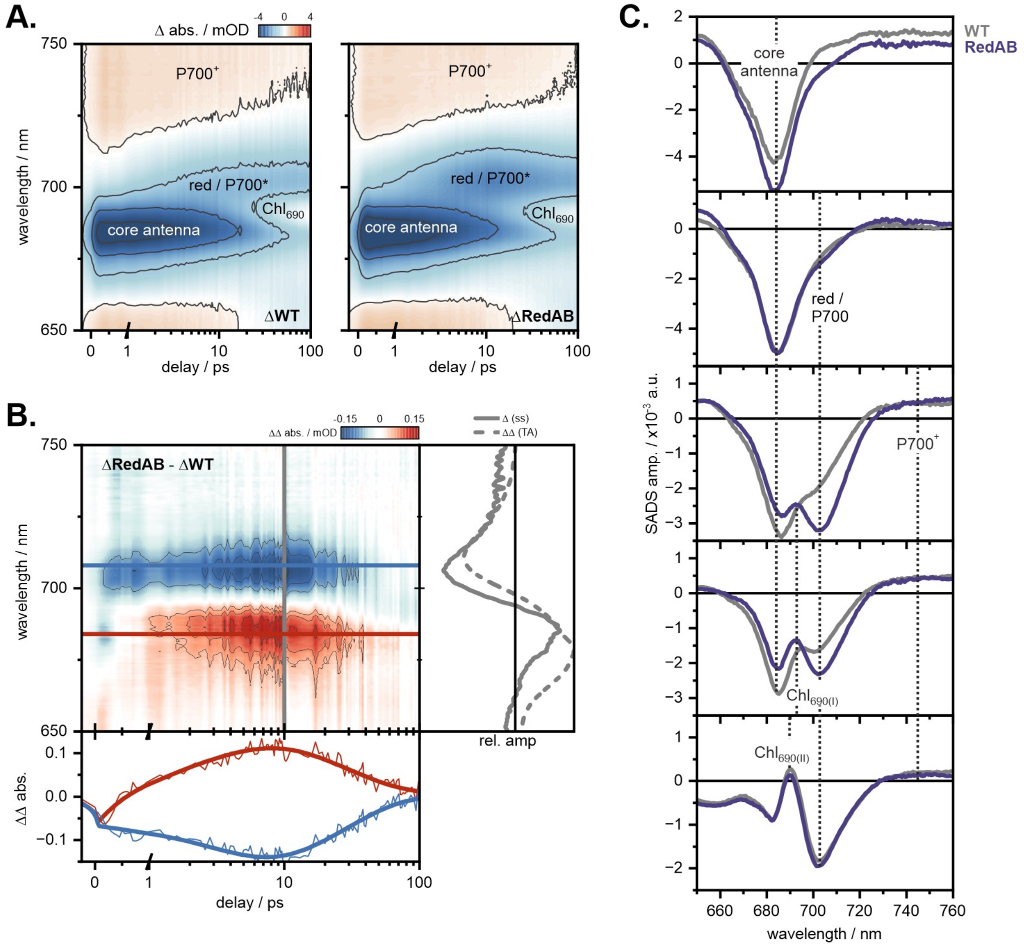

**Figure s14: A.** Ultrafast transient data of WT and RedAB**. A.** Normalized TA data measured on wt6803 and RedAB purified PSI trimers**. B.** Double difference (ΔΔ) spectra of RedAB after subtraction of the ΔWT data. The spectral cuts (grey, solid line) are overlayed with the inverted and scaled steady state difference (Δ) spectra (grey, dashed lines). The temporal evolution of the ΔΔ data is extracted by horizontal cuts in the core antenna ΔΔ band (red) and red/P700* ΔΔ band (blue). **C.** Overlayed SADS of WT (grey) and RedAB (purple) obtained by GTA.

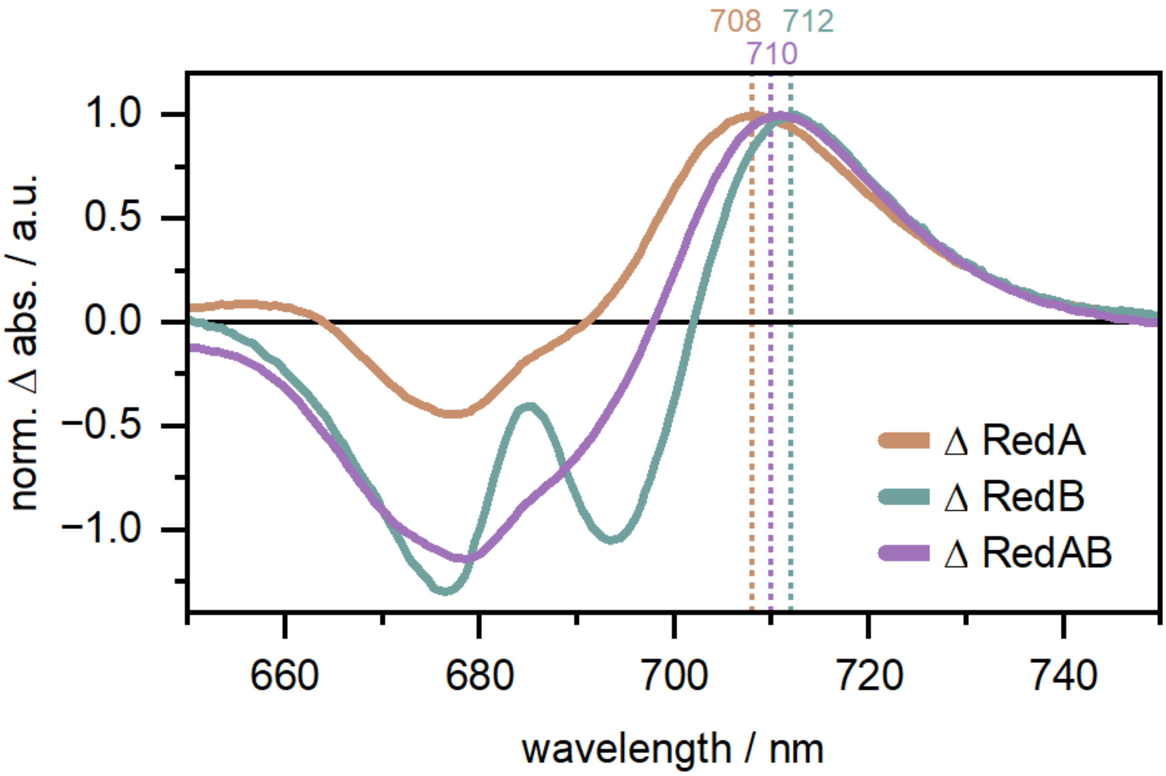

**Figure s15.** Steady-state difference spectra of each mutant after subtraction of the wildtype spectrum. The RedB mutant shows a significant relative reduction of the antenna band absorption due to its proximity to the bulk antenna network. The induced red shifts peak at 708 nm (RedA), 710 nm (RedAB) and 712 nm (RedB).

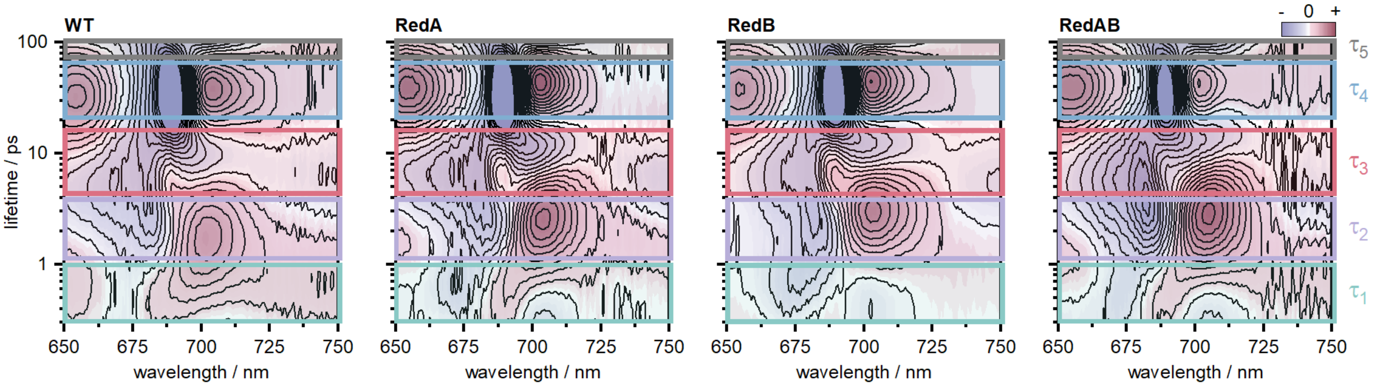

**Figure s16.** Lifetime density maps (LDMs) of each variant. Every LDM is composed of five similar lifetime component ranges τ1-τ5. The initial decay of the chl-associated ESA signatures in the blue and red edge of the spectral window and the initial decay of the main core antenna band at ~680 nm occurs within < 1 ps (τ1). This is followed by a red shift of the GSB from ~680 nm to ~700 nm at 2-3 ps corresponding to the population of P700* which is slightly delayed in all variants compared to the wildtype (in the order WT < RedB < RedAB < RedA). The red shoulder ($\lambda_{max}$ ~ 705-715 nm) of this transiently formed GSB increases further in the red variants and is therefore indicative of absorption contributions by the additional red chls. It is important to note, that the wildtype also contains red absorbing chls which contribute to this spectral region. The population of the red absorbing sites is temporally stretched (especially in RedA and RedAB) and ranges from 4-10 ps (see also **figure s17**). The rise of difference absorption above 725 nm, which is associated with the formation of the P700+ radical, occurs on a similar timescale. The absorption at ~695 nm, indicative for the neutral $Chl_{690}$, grows within < 10 ps and is strongly pronounced in RedB. The final transient spectrum which is associated with the $P700^{+}/A_1$ species is then formed within 30-40 ps (τ4) and remains constant in the infinity lifetime component (τ5).

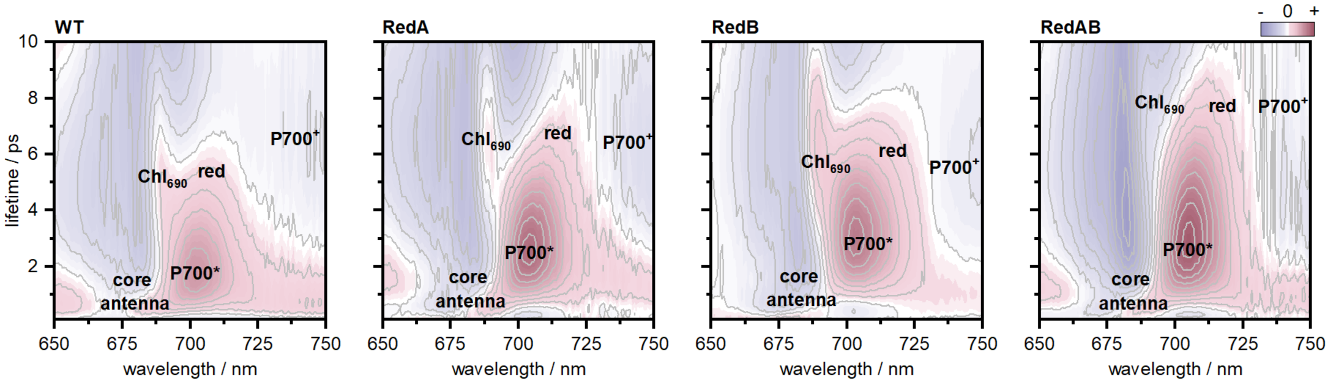

**Figure s17.** Lifetime density maps (LDMs) of each variant in linear scaling to enhance comparability. This temporal range is mostly dominated by the equilibration dynamics between the core antenna, P700* and red. The red population accumulates up to the 4-10 ps window. The transition to the charge separated species is indicated by the $Chl_{690}$ and $P700^{+}$ signatures.

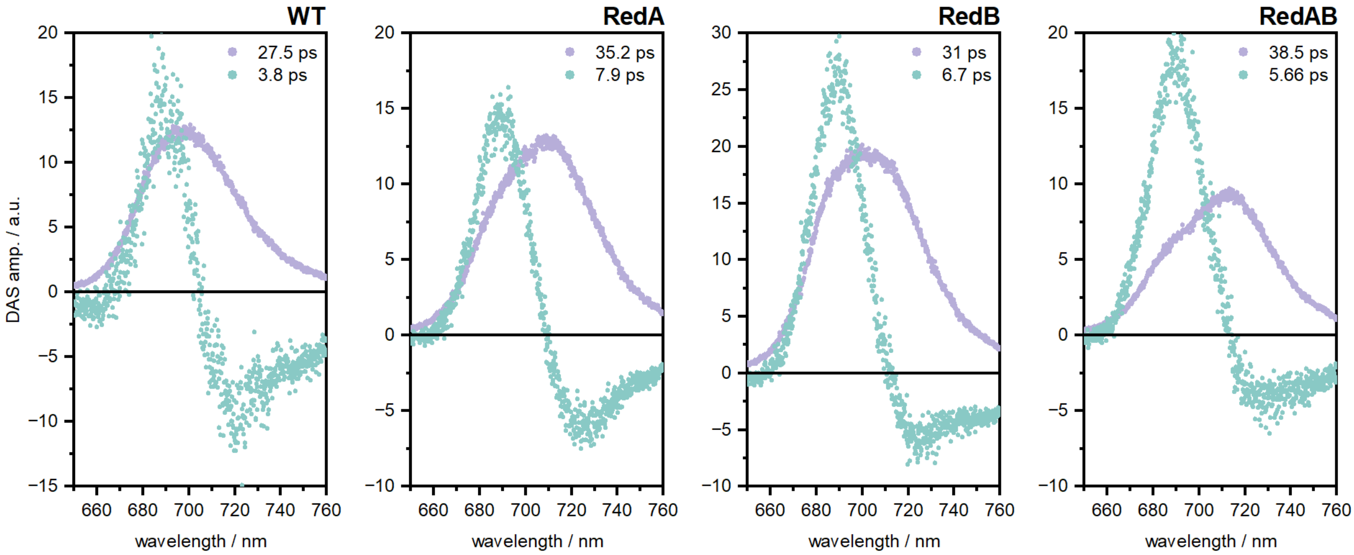

**Figure s18.** Decay-associated spectra (DAS) obtained from global analysis of the time-resolved fluorescence measurements. The slow component (purple) is associated with the trapping step. The fast component (green) is usually solely assigned to the energy equilibration, however as inferred from the fs-TA data, this temporal range is also coincident with the charge separation.

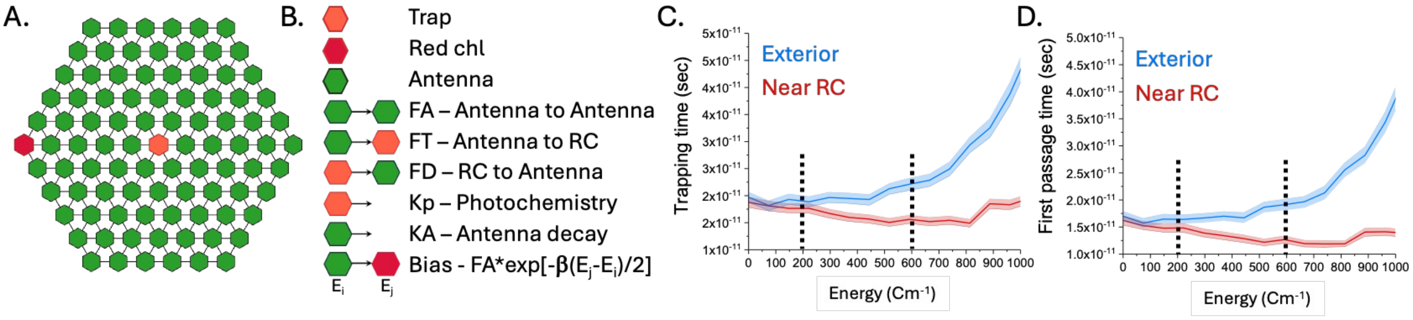

**Figure s19: The effect of red site location on trapping time in an idealized antennae system. A**. The array used to test different red sites location is used.. An array exterior ‘A’, or a RC proximal ‘B’, red chl location. **B**. Excited state migration was modeled as a random walk on these latices using 6 rates. All Antennae nodes (green hexagons) are treated iso-energetically, while distinct rates or biases are introduced between the central node (named RC, which also has a photochemical rate, Kp, associated with it), a location assigned to a red chl, and the bulk antenna. **C**. and **D**. Simulating both red site location scenarios over a range of energy values (kt at 25 $^{o}$C and the estimated energetic gap for both the RedA and RedB mutants are highlighted using dotted lines). Plot ‘C’ shows the effects on trapping time (M0), and ‘D’ the effect on the first passage time (the time it takes the excited state to reach the RC for the first time). It’s clear that placing a red chl at peripheral location increase both values compared to a RC proximal scenario. Confidence intervales (shaded) represent ±5 % bootstrapping values. The rates used in the simulation were: FA = $10^{13}$ $sec^{-1}$, FT = $10^{12}$ $sec^{-1}$, FD = $1.85*10^{12}$ $sec^{-1}$, KP = $10^{12}$ $sec^{-1}$, KA = 0 $sec^{-1}$.

### Table s1. Data collection and refinement statistics

| | Monomer (EMD-71005, PDB 9OY9) | Trimer (EMD-71048, PDB 9OZ3) |
|---|---|---|
| **Data collection and processing** | | |
| Magnification | 165,000 | |
| Voltage (kV) | 300 | |
| Electron exposure ($e^{-}/Å^{2}$) | 45 | |
| Defocus range (μm) | 0.5 - 3 | |
| Pixel size (Å) | 0.731 | |
| Symmetry imposed | C3 expanded | C1 |
| Initial particle images (no.) | 1,508,670 | |
| Final particle images* (no.) | 857,484 | 285,828 |
| Map resolution (Å) | 1.76 | 1.84 |
| FSC threshold | 0.143 | |
| **Refinement** | | |
| Initial model used (PDB code) | 6UZV | |
| Model resolution (Å) | 1.75/1.84 | 1.88/1.99 |
| FSC threshold | 0.143,0.5 | |
| Map sharpening *B* factor ($Å^{2}$) | 0 | 0 |
| Model composition | | |
| Nonhydrogen atoms | 25,586 | 76,301 |
| Protein residues | 2241 | 6729 |
| Ligands | 141 | 436 |
| *B* factors ($Å^{2}$) | | |
| Protein | 47.39 | 55.33 |
| Ligand | 51.46 | 60.11 |
| Water | 49.50 | 57.50 |
| R.m.s. deviations | | |
| Bond lengths (Å) | 0.007 | 0.004 |
| Bond angles (°) | 0.943 | 0.868 |
| **Validation** | | |
| MolProbity score | 1.17 | 0.96 |
| Clashscore | 2.39 | 1.95 |
| Poor rotamers (%) | 0.44 | 0.35 |
| Ramachandran plot | | |
| Favored (%) | 97.16 | 98.18 |
| Allowed (%) | 2.75 | 1.82 |
| Disallowed (%) | 0.09 | 0 |

### Table s2. Coupling values for RedA chls.

| | B31-B32 | B31-B33 | B32-B33 |
|---|---|---|---|
| 6803 - 5oy0 | -75.91 | | |
| TV – 1JB0 | -56.49 | -27.64 | -32.81 |
| RedA – 6uzv | -62.08 | -28.05 | -52.22 |
| RedAB | -84.89 | -29.32 | -62.03 |

### Table s3. Coupling values for RedB chls.

| | B7-A31 | B7-A32 | A31-A32 |
|---|---|---|---|
| 6803 - 5oy0 | -44.32 | 118.40 | 78.21 |
| TV – 1JB0 | -40.99 | 130.41 | 81.94 |
| RedA – 6uzv | -38.90 | 118.02 | 90.30 |
| RedAB | -41.38 | 123.15 | 82.53 |

**Table s4.** Lifetime ranges obtained by LDA (figure s16).

| | $\tau_1$ / ps | $\tau_2$ / ps | $\tau_3$ / ps | $\tau_4$ / ps | $\tau_5$ / ps |
|---|---|---|---|---|---|
| | 0.3 - 1 | 1 - 5 | 5 - 18 | 20 - 50 | inf. |

**Table s5.** Lifetimes obtained by exponential fitting of the $\Delta\Delta$ datasets shown in figures 4 A+B and s14 B.

| | $\tau_1$ / ps | $\tau_2$ / ps | $\tau_3$ / ps | $\tau_4$ / ps |
|---|---|---|---|---|
| **RedA** | 0.7 | 4.5 | 27 | inf. |
| **RedB** | 0.7 | 3.3 | 22 | inf. |
| **RedAB** | 0.7 | 3.6 | 28 | inf. |

**Table s6. Lifetimes corresponding to the proposed 5-compartment model within the TGA**. The lifetimes were soft-fixed within a margin of 5-10% based on the values acquired in S5 and the time-resolved fluorescence data.

| | **$\tau_1$ / ps** | **$\tau_2$ / ps** | **$\tau_3$ / ps** | **$\tau_4$ / ps** | **$\tau_5$ / ps** |
|---|---|---|---|---|---|
| **WT** | 0.7 | 3.0 | 8.5 | 30 | inf. |
| **RedA** | 0.7 | 4.6 | 9 | 35 | inf. |
| **RedB** | 0.7 | 3.4 | 7.2 | 32 | inf. |
| **RedAB** | 0.7 | 3.85 | 12 | 38 | inf. |
